# A domesticated prophage enzyme links bacterial metabolism, episymbiosis and phage fitness

**DOI:** 10.64898/2026.10.08.757619

**Authors:** Nusrat Nahar, Ajay Kumar, Alex S. Grossman, Jeffrey S. McLean, Batbileg Bor, Pu-Ting Dong, Xuesong He

## Abstract

Prophages are increasingly recognized as drivers of bacterial adaptation, yet how domesticated prophage genes shape microbial symbiosis remains poorly understood. Here, we identify GDPD_Xhp1_, a glycerophosphodiester phosphodiesterase encoded by prophage Xhp1 of *Schaalia odontolytica* XH001, as a metabolic regulator linking bacterial physiology, episymbiosis and phage fitness. Association with the epibiotic bacterium *Nanosynbacter lyticus* TM7x strongly induced *gdpd*_Xhp1_ expression and triggered GDPD_Xhp1_-dependent lipid droplet accumulation in XH001. GDPD_Xhp1_ orchestrated glycerophospholipid remodeling, membrane biophysical properties, respiration and cell-surface glycan composition in XH001, thereby promoting TM7x episymbiosis. Remarkably, GDPD_Xhp1_ was also required for productive Xhp1 infection by remodeling susceptible bacterial host envelope structures to facilitate phage–host interactions. Together, our findings reveal a domesticated prophage enzyme that contributes to bacterial metabolism, interspecies symbiosis and viral reproduction, highlighting prophage as an overlooked source of metabolic innovation in microbial symbiosis.

## Introduction

Bacteriophages shape bacterial evolution not only through predation and horizontal gene transfer but also through the long-term retention of prophage genes within bacterial genomes^1–3^. Increasing evidence suggests that some prophage genes become domesticated and repurposed for host-beneficial functions, contributing to stress adaptation, virulence, and metabolism^4–7^. However, how domesticated prophage genes influence complex interspecies interactions remains poorly understood.

The oral bacterium *Schaalia odontolytica* XH001 (hereafter XH001) forms an obligate episymbiotic association with the Patescibacteriota, also known as Candidate Phyla Radiation (CPR), bacterium *Nanosynbacter lyticus* TM7x (hereafter TM7x)^8–12^. TM7x grows on the surface of XH001 and induces extensive physiological remodeling of its host, including altered gene expression^13, 14^, membrane fluidity change^15^, and the accumulation of intracellular lipid droplets (LDs)^15^. Although LDs formation is one of the most phenotypically striking hallmarks of TM7x episymbiosis^15^, the mechanisms linking epibiont colonization to host metabolic remodeling remain unknown.

Glycerophosphodiester phosphodiesterases (GDPDs) hydrolyze glycerophosphodiesters to generate glycerol-3-phosphate (G3P)^16^, a key intermediate connecting phospholipid turnover, energy metabolism, and membrane homeostasis^17, 18^. While GDPDs are widespread among bacteria, related enzymes are also encoded by numerous bacteriophages^19–29^, where they may facilitate infection through modification of the bacterial cell envelope^30^. Whether phage-derived GDPDs can be co-opted to regulate bacterial metabolism or symbiotic interactions has not been explored.

In addition to serving as a bacterial host for TM7x, XH001 harbors an intact inducible prophage Xhp1 capable of entering a productive lytic cycle^31, 32^. Here we show that Xhp1-encoded GDPD, named GDPD_Xhp1_, functions as a domesticated metabolic regulator that links bacterial physiology, episymbiosis, and phage fitness. We demonstrate that GDPD_Xhp1_ is required for XH001 glycerophospholipid remodeling, membrane homeostasis, and efficient TM7x colonization. TM7x episymbiosis induces overexpression of *gdpd*_Xhp1_ and results in enhanced LDs biogenesis. Meanwhile, the same enzyme is also required for productive infection by the resident prophage Xhp1. Our findings reveal a domesticated prophage enzyme that links bacterial metabolism, interspecies symbiosis, and viral fitness, highlighting prophages as an underappreciated source of metabolic innovation in microbial interactions.

## Results

### TM7x-induced LD formation in XH001 requires the prophage Xhp1-encoded GDPD_Xhp1_

To investigate whether prophage domestication contributes to the metabolic remodeling associated with episymbiosis, we first examined the role of the resident prophage Xhp1 in TM7x-induced LD biogenesis. Because association with TM7x is known to stimulate robust LD accumulation in XH001^15^, we stained neutral lipids with LipidSpot and compared LD formation in wild-type (WT) XH001 and the prophage-cured derivative XH001ΔXhp1 grown either in monoculture or in association with TM7x. Whereas WT XH001 exhibited extensive LD accumulation during TM7x association (Fig. 1A), the prophage-cured strain failed to induce LD formation in response to TM7x (Fig. 1B). Both XH001 and XH001ΔXhp1 exhibited only low basal levels of LD under monoculture conditions (Fig. 1C, D). Quantitation of single-LD fluorescence intensity confirmed the visual observation and revealed the strongest staining signal in TM7x-associated XH001 compared with other groups. Meanwhile, the basal level of LD was significantly lower in XH001ΔXhp1 relative to WT under monoculture conditions (Fig.1E).

**Figure 1:**
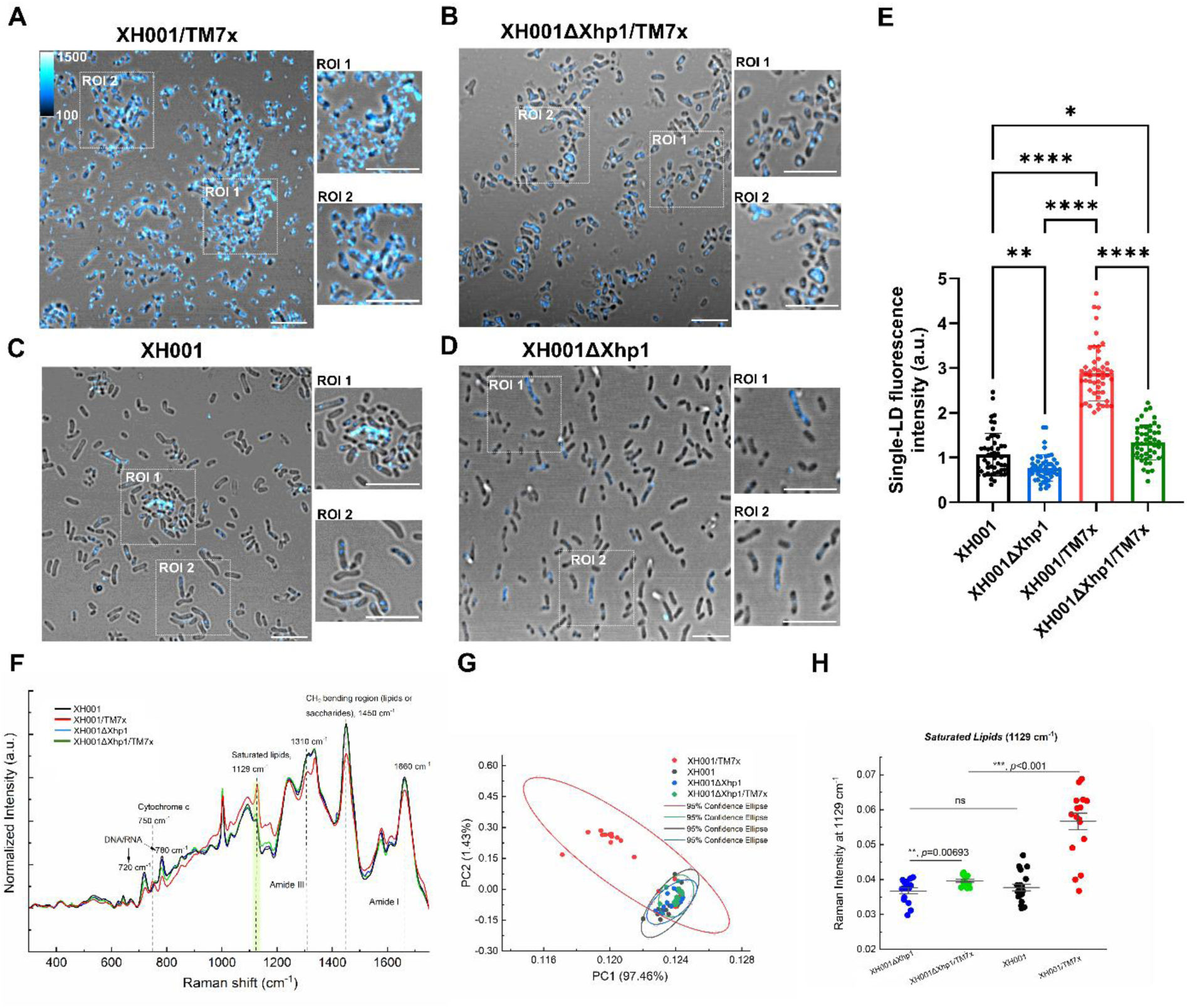
Prophage Xhp1 is required for TM7x-induced LD formation. (A-D) Representative confocal laser scanning microscopy (CLSM) images of LipidSpot-stained XH001 and XH001ΔXhp1 either in monoculture or in association with TM7x. Scale bar = 5 μm. (E) Quantification of single-LD fluorescence intensity in all strains. Data were collected from at least 50 individual lipid droplets from three biological replicates per group. Statistical significance was determined by one-way ANOVA followed by Tukey’s multiple comparison test; *P* < 0.05 (*), *P* < 0.01 (**), *P* < 0.0001 (****). (F) Raman spectra of XH001, XH001ΔXhp1, XH001/TM7x, and XH001ΔXhp1/TM7x from three biological replicates following vector normalization. Selected Raman peaks are indicated by dashed lines and shaded box. (G) PCA analysis of Raman spectra from all experimental groups; each dot represents an individual Raman spectrum. (H) Quantification and statistical analysis of Raman intensity at 1128 cm^−1^, corresponding to saturated lipids. Statistical significance was determined using a two-tailed unpaired Student’s *t*-test;*P* < 0.01 (**), *P* < 0.001 (***), ns (not significant).

To further characterize the metabolic differences associated with TM7x-induced LD accumulation, we performed label-free Raman spectroscopy. The Raman fingerprint region spanning 300–1800 cm^−1^ captures vibrational signatures associated with major cellular biomolecules^33^ and revealed distinct spectral profiles among the experimental groups (Fig. 1F). Principal component analysis (PCA) of the Raman spectra showed clear separation of TM7x-associated XH001 from the other groups (Fig. 1G). Consistent with the increased LD accumulation observed by LipidSpot staining and with the Raman intensity previously reported by Dong *et al*^15^, XH001/TM7x exhibited a significantly higher Raman intensity associated with saturated lipid content compared with the other groups (*P* < 0.001) (Fig. 1H). Together, these findings demonstrate that the prophage Xhp1 is required for TM7x-induced LD accumulation and suggest that one or more prophage-encoded functions contribute to lipid remodeling.

To identify the prophage determinant responsible for TM7x-induced lipid remodeling, we examined the Xhp1 genome and identified a prophage gene, APY09_09275, predicted to encode a glycerophosphodiester phosphodiesterase (GDPD). Sequence analyses revealed that the encoded protein GDPD_Xhp1_ was distantly related to previously characterized bacterial GDPDs, sharing 19.1% amino-acid identity with *Escherichia coli* UgpQ^34^ and 18.3% identity with *Thermoanaerobacter tengcongensis* ttGDPD^35^ (Supplementary Fig. 1). Despite this sequence divergence, conserved-domain analysis and structural modeling using NCBI Conserved Domain Search database (CD-Search) identified a canonical GDPD architecture consisting of an N-terminal extension and a C-terminal phospholipase C-like phosphodiesterase domain containing the conserved metal-coordinating residues characteristic of the UgpQ superfamily (Fig. 2A). Consistent with these predictions, purified recombinant GDPD_Xhp1_ hydrolyzed glycerophosphocholine (GPC) to release G3P (Fig. 2B), confirming that APY09_09275 encodes a functional GDPD.

**Figure 2:**
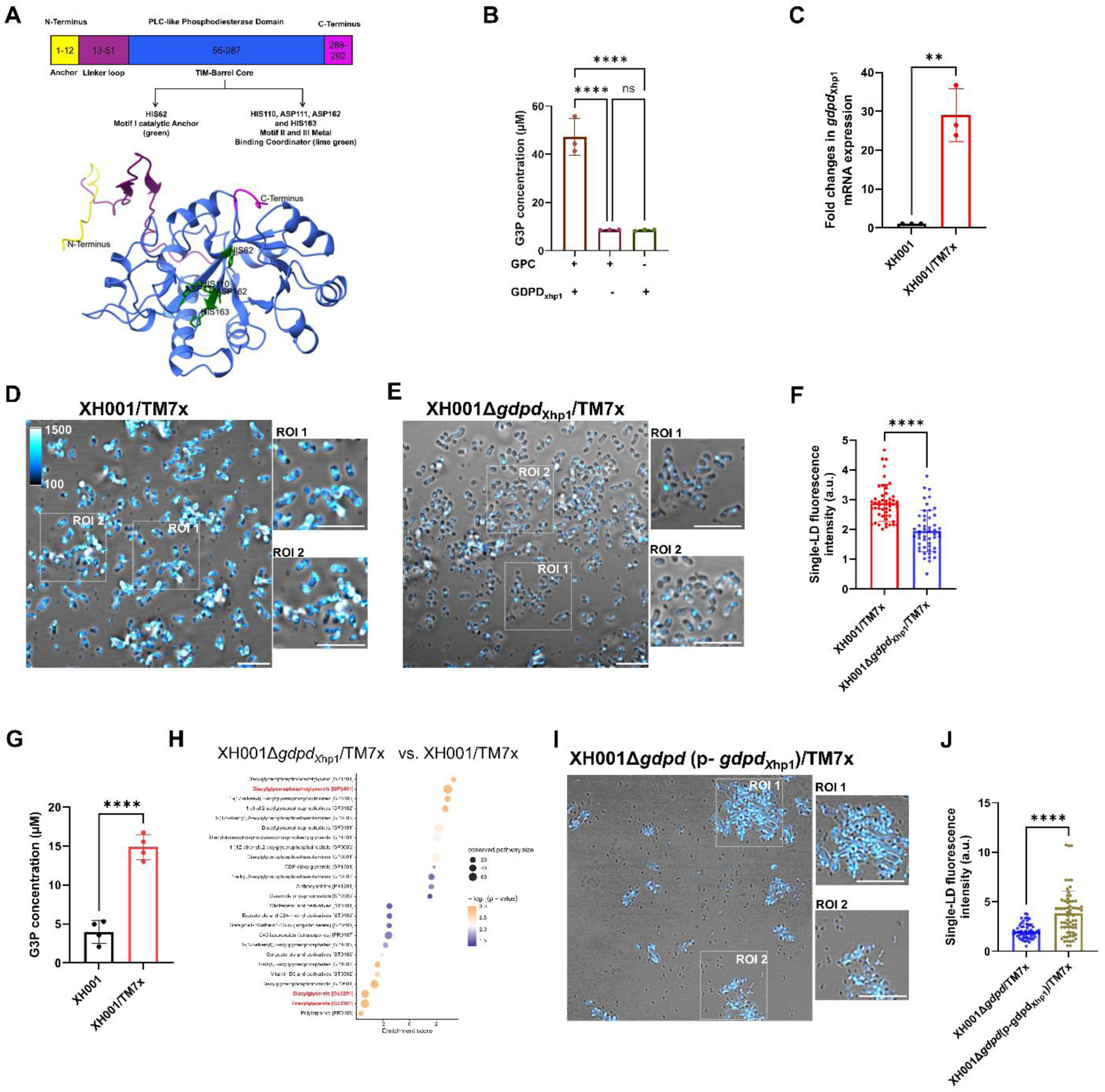
GDPD_Xhp1_ is a functional prophage-encoded glycerophosphodiester phosphodiesterases that drives TM7x-induced LD biogenesis. (A) Predicted structure of the 293-amino acid GDPD_Xhp1_ showing the N-terminal region and the conserved PLC-like phosphodiesterase domain containing the predicted catalytic residues. (B) Enzymatic activity of purified His-tagged GDPD_Xhp1_ toward glycerophosphocholine (GPC) *in vitro*, measured by G3P production. Statistical significance was determined by one-way ANOVA followed by Tukey’s multiple comparison test; *P* < 0.0001 (****), ns = not significant. (C) Relative expression of *gdpd*_Xhp1_ in XH001 monoculture and coculture, demonstrating increased *gdpd*_Xhp1_ expression during TM7x symbiosis. Statistical significance was determined by unpaired Student’s *t* test. *P* < 0. 01 (**). (D-E) Representative confocal laser scanning microscopy (CLSM) images of LipidSpot-stained XH001/TM7x (D) and XH001Δ*gdpd*_Xhp1_/TM7x (E) cocultures. Scale bar = 5 μm. (F) Quantification of single-LD fluorescence intensity in all strains. Data were collected from at least 50 individual lipid droplets from three biological replicates per group. Statistical significance was determined by Welch’s *t-*test; *P* < 0.0001 (****). (G) Quantification of intracellular G3P levels showing increased G3P abundance in XH001/TM7x cocultures compared with monoculture. Statistical significance was determined by Welch’s *t-*test; *P* < 0.0001 (****). (H) Lipid class enrichment analysis of XH001Δ*gdpd*_Xhp1_/TM7x relative to XH001/TM7x. Enrichment scores indicate whether lipid classes are overrepresented among metabolites increased (positive values) or decreased (negative values) in the Δ*gdpd*_Xhp1_ mutant. Triacylglycerols [GL0301], diacylglycerols [GL0201] and diacylglycerophosphoglycerols [GL0401] are highlighted in red. (I) Representative confocal laser scanning microscopy (CLSM) image of the complemented strain XH001Δ*gdpd*(p-*gdpd*_Xhp1_)/TM7x stained with LipidSpot showing restoration of LD formation. (J) Quantification of single-LD fluorescence intensity in XH001Δ*gdpd*_Xhp1_/TM7x and the complemented strain XH001Δ*gdpd*(p-*gdpd*_Xhp1_)/TM7x cocultures. Statistical significance was determined by Welch’s *t-*test; *P* < 0.0001 _(****)._

We next examined whether *gdpd*_Xhp1_ expression was associated with TM7x-induced lipid remodeling. Quantitative RT–PCR revealed that TM7x association induced *gdpd*_Xhp1_ expression more than 20-fold relative to XH001 monocultures (Fig. 2C). To determine whether GDPD_Xhp1_ contributes to TM7x-dependent LD biogenesis, we generated a XH001Δ*gdpd*_Xhp1_ mutant. Whereas WT XH001 accumulated abundant LDs during TM7x association, deletion of *gdpd*_Xhp1_ nearly eliminated LD formation, phenocopying the prophage-cured strain (Fig. 2D, E, F and Fig. 1B). Together, these results identify GDPD_Xhp1_ as the prophage-encoded factor required for TM7x-induced lipid remodeling and establish a direct link between prophage domestication and host metabolic reprogramming during episymbiosis.

### GDPD_Xhp1_ drives neutral lipid accumulation and LD biogenesis in XH001 during symbiosis with TM7x

To investigate the metabolic consequences of GDPD_Xhp1_ activity, we first quantified intracellular G3P, the product of glycerophosphodiester hydrolysis. XH001/TM7x symbiotic cocultures contained significantly higher levels of G3P than XH001 monocultures (*P* < 0.0001; Fig. 2G), consistent with enhanced LD accumulation in XH001 during its episymbiosis with TM7x, as G3P is the precursor for LD formation^15^. We next examined the effects of *gdpd*_Xhp1_ deletion on cellular lipid composition using untargeted lipidomics. PCA analysis revealed clear separation between XH001/TM7x and XH001Δ*gdpd*_Xhp1_/TM7x samples (Supplementary Fig. 2A), indicating extensive metabolic remodeling in the absence of GDPD_Xhp1_. In total, 321 metabolites differed significantly between the two groups (*P* < 0.05; Supplementary Fig. 2B), spanning 26 lipid classes and metabolic pathways (Fig. 2H).

The most pronounced changes involved neutral storage lipids. WT XH001/TM7x cocultures, which accumulated abundant LDs, were enriched in triacylglycerols (GL0301) and diacylglycerols (GL0201), whereas both lipid classes were markedly depleted in XH001Δ*gdpd*_Xhp1_/TM7x cocultures (Supplementary Fig. 2C, D). Conversely, the mutant retained higher levels of membrane-associated glycerophospholipids (GP0401), consistent with a failure to redirect membrane lipid pools toward neutral lipid storage. These findings indicate that GDPD_Xhp1_ is a key metabolic node linking glycerophospholipid turnover to LD formation and suggest that TM7x-induced lipid remodeling proceeds through the conversion of membrane lipids into neutral storage lipids characteristic of actinobacterial lipid droplets^36^.

To determine whether GDPD_Xhp1_ is sufficient to drive this process, we expressed *gdpd*_Xhp1_ in the deletion mutant (XH001Δ*gdpd*_Xhp1_). Plasmid-borne expression of *gdpd* (p-*gdpd*_Xhp1_) restored LD formation, promoting lipid accumulation in the deletion mutant, with or without TM7x present (Figure 2I, Supplementary Fig.3). Consistent with these observations, quantification of single-LD fluorescence intensity revealed a significantly greater abundance of LDs in the complemented XH001Δ*gdpd*(p-*gdpd*_Xhp1_)/TM7x coculture than in the XH001Δ*gdpd*_Xhp1_*/*TM7x coculture (*P* < 0.0001, Figure 2J). Together, these results demonstrate that GDPD_Xhp1_ is necessary for LD biogenesis and identify a prophage-derived enzyme as the molecular driver of TM7x-induced lipid remodeling in XH001.

**Figure 3.**
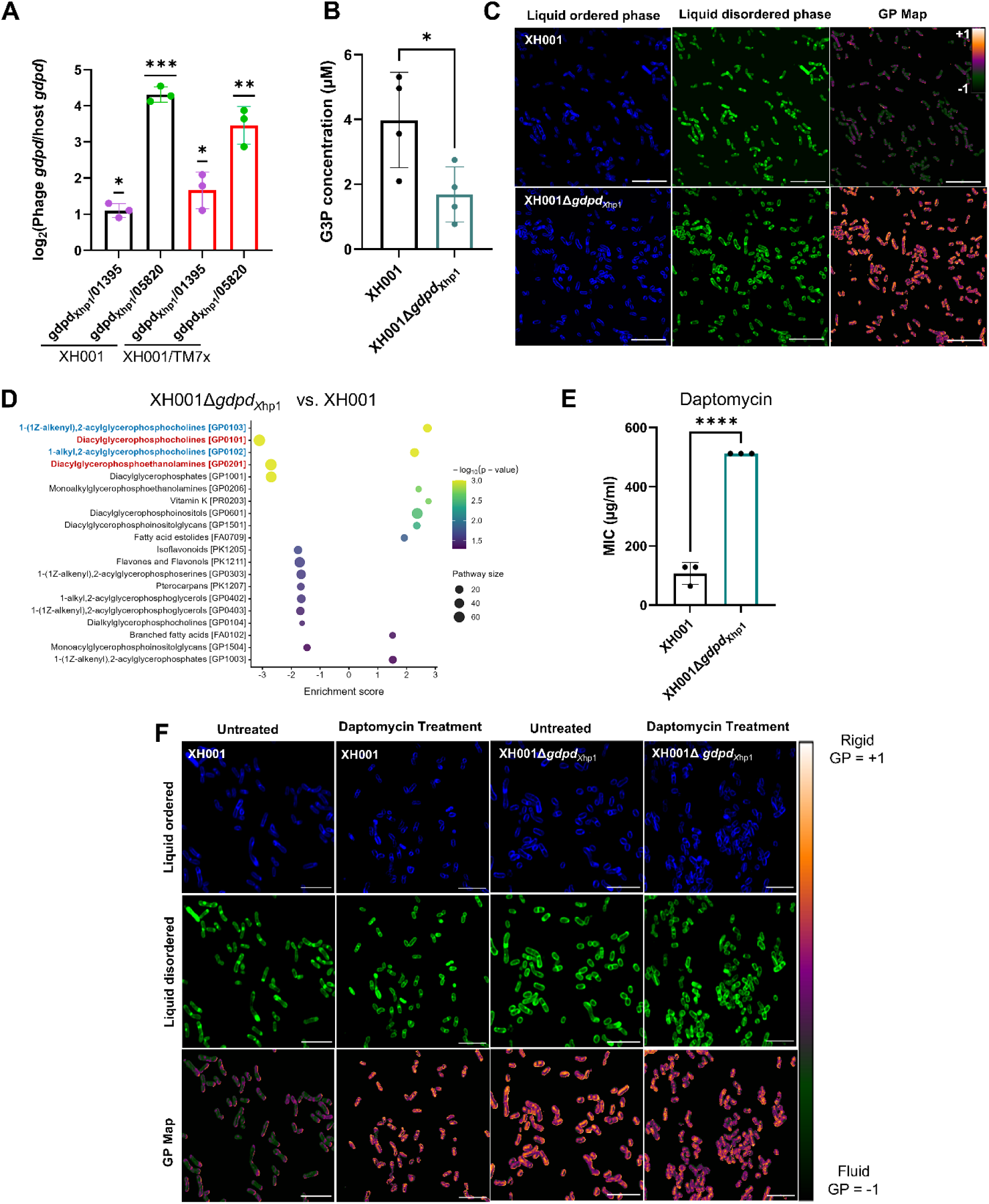
GDPD_Xhp1_ contributes to membrane fluidity in XH001. (A) Relative expression of prophage-encoded *gdpd*_Xhp1_ and chromosomally encoded GDPD homologues (APY09_01395 and APY09_05820) in XH001 under TM7x-free and TM7x-associated conditions. Expression is shown as the ratio of *gdpd*_Xhp1_ to each chromosomal homologue. Statistical significance was determined using a one-sample *t*-test on log₂-transformed expression ratios against a theoretical mean of 0 (equivalent to a ratio of 1). Data are presented as mean ± SD. *P* < 0.05 (*), *P* < 0.01 (**), *P* < 0.001 (***). (B) Intracellular G3P levels in WT XH001 and Δ*gdpd*_Xhp1_ mutant. Statistical significance was determined by Welch’s *t-*test; *P* < 0.05 (*). (C) Laurdan fluorescence imaging of WT XH001 and XH001Δ*gdpd*_Xhp1_. Images show liquid-ordered phase, liquid-disordered phase, and generalized polarization (GP) maps. GP values range from −1 (more fluid) to +1 (more rigid). Scale bar = 5 μm. (D) Lipid class enrichment analysis of XH001Δ*gdpd*_Xhp1_ relative to XH001. Enrichment scores indicate whether lipid classes are overrepresented among metabolites increased (positive values) or decreased (negative values) in the Δ*gdpd*_Xhp1_ mutant. (E) MIC analysis of WT XH001 and XH001Δ*gdpd*_Xhp1_ strains against daptomycin. Statistical significance was determined by two-tailed unpaired Student’s *t-*test; *P* < 0.0001 (****). (F) Laurdan-based fluorescence imaging of WT XH001 and XH001Δ*gdpd*_Xhp1_ strains following 30 min daptomycin exposure. Daptomycin increased membrane rigidity in XH001 strain but caused no detectable GP shift in the Δ*gdpd*_Xhp1_ mutant. Scale bar = 5 μm.

### GDPD_Xhp1_ contributes to membrane homeostasis and antibiotic susceptibility

G3P serves as a key precursor for both triacylglycerol synthesis in LDs and membrane glycerolipid biogenesis^37^. To determine the contribution of GDPD_Xhp1_ to glycerophosphodiester metabolism in XH001, we first compared the expression of the three *gdpd* homologs encoded by XH001. Although XH001 carries two chromosomal *gdpd* homologs (APY09_01395 and APY09_05820), *gdpd*_Xhp1_ was expressed at significantly higher levels under both monoculture and TM7x-associated conditions (Fig. 3A), identifying the prophage-derived enzyme as the predominant GDPD in this organism. Consistent with this result, deletion of *gdpd*_Xhp1_ significantly decreased intracellular G3P abundance (*P* < 0.05; Fig. 3B), indicating that prophage-encoded GDPD_Xhp1_ is the principal contributor to glycerophosphodiester turnover and G3P production in XH001.

Because glycerophospholipid metabolism is intimately linked to membrane organization^38^, we next assessed membrane fluidity using Airyscan super-resolution microscopy combined with Laurdan membrane imaging^15, 39^. Compared with WT XH001, XH001Δ*gdpd*_Xhp1_ displayed a marked increase in generalized polarization (GP) values, indicative of increased membrane order and reduced membrane fluidity (Fig. 3C). These findings suggest that GDPD_Xhp1_-dependent glycerolipid metabolism contributes to the biophysical properties of the XH001 membrane.

To define the lipidomic basis of this phenotype, we performed untargeted lipidomics on WT XH001 and XH001Δ*gdpd*_Xhp1_ monocultures. PCA revealed clear segregation of the two strains (Supplementary Fig. 4A), indicating extensive lipidome remodeling following loss of GDPD_Xhp1_. In total, 50 annotated metabolites and 18 lipid classes were significantly altered in the mutant (Fig. 3D and Supplementary Fig. 4B). The most pronounced changes occurred within phosphatidylcholine metabolism, characterized by depletion of diacylglycerophosphocholines and concomitant enrichment of ether-linked phosphatidylcholine species, including plasmalogen-like and alkyl-acyl derivatives (Fig. 3D and Supplementary Fig. 4C-E). The accumulation of ether-linked phospholipids is consistent with the increased membrane order observed in the Δ*gdpd*_Xhp1_ mutant^40, 41^.

Given the changes in membrane composition and biophysical properties, we next examined the antimicrobial susceptibility of XH001 and XH001Δ*gdpd*_Xhp1_ to membrane-targeting antibiotics such as daptomycin. XH001Δ*gdpd*_Xhp1_ exhibited increased resistance to daptomycin relative to WT cells (Fig. 3E). To determine whether this phenotype was associated with altered membrane dynamics, we evaluated the membrane fluidity through Laurdan imaging following antibiotic exposure. Daptomycin treatment increased GP values in WT cells, consistent with increased membrane order^42^, whereas GP values remained unchanged in the Δ*gdpd*_Xhp1_ mutant (Fig. 3F). These findings indicate that loss of GDPD_Xhp1_ alters both basal membrane organization and the membrane response to antibiotic challenge. Collectively, these results identify GDPD_Xhp1_ as a major determinant of phospholipid homeostasis in XH001. By promoting glycerophosphodiester catabolism and G3P production, the prophage-derived enzyme influences membrane lipid composition, membrane order, and susceptibility to a membrane-targeting antibiotic. These observations reveal an unexpected contribution of a domesticated prophage gene to the maintenance of bacterial membrane physiology.

### GDPD_Xhp1_ links central metabolism to envelope remodeling and episymbiotic colonization

Because G3P lies at the intersection of phospholipid metabolism and cellular energy pathways^17, 18^, we examined the global consequences of GDPD_Xhp1_ loss on XH001 physiology. Comparative transcriptomic analysis revealed extensive transcriptional reprogramming in the Δ*gdpd*_Xhp1_ mutant, with prominent changes in genes involved in respiration and central metabolism (Fig. 4A). In particular, operons encoding NADH:quinone oxidoreductase and nitrate reductase were significantly upregulated relative to WT XH001, indicating altered redox and energy metabolism following disruption of glycerophosphodiester turnover.

**Figure 4.**
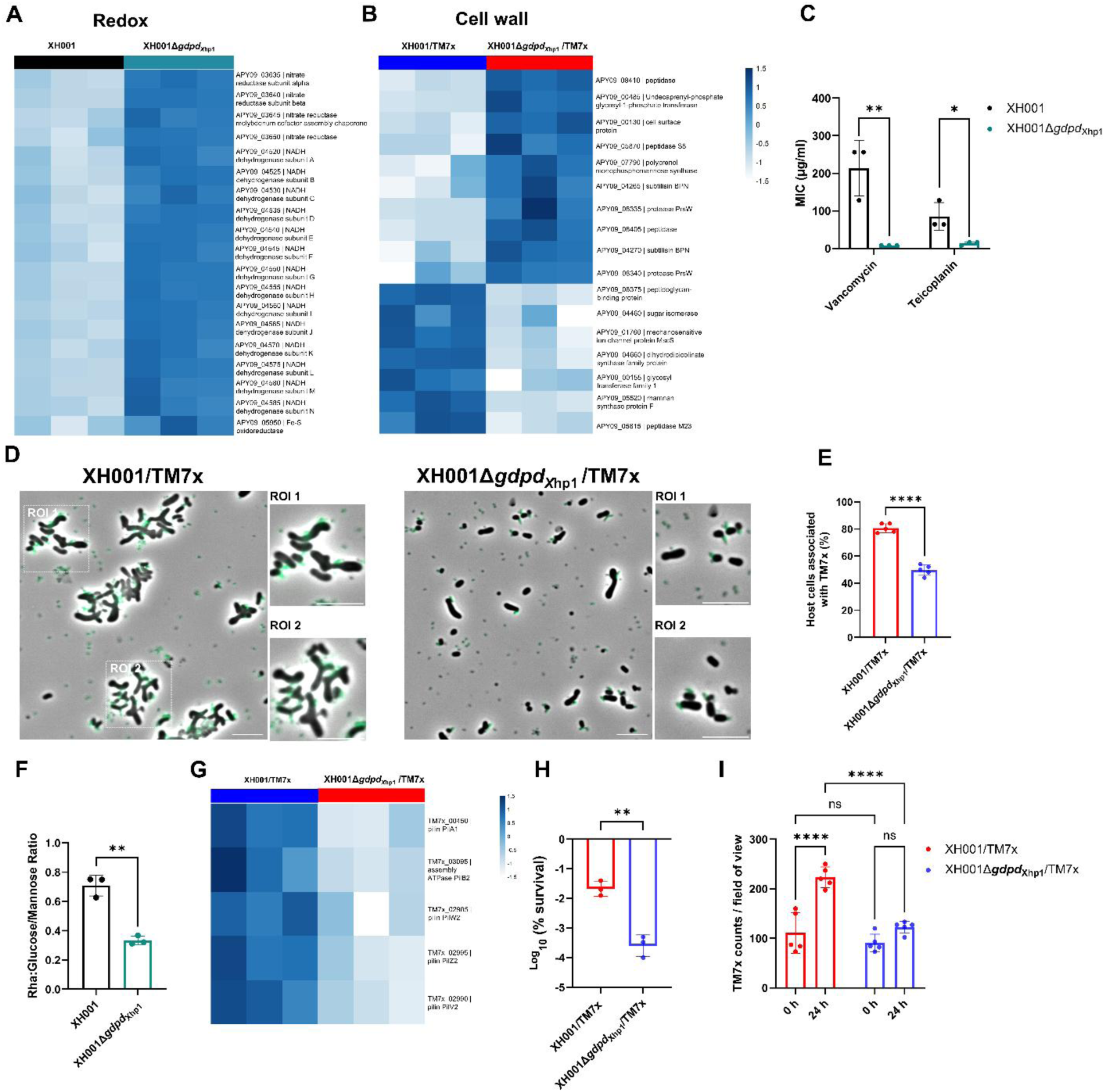
GDPD_Xhp1_ affects host surface architecture, TM7x association and symbionts persistence. (A) RNA-seq heatmap highlighting differential expression of genes associated with NADH-quinone oxidoreductase and nitrate reductase pathways in XH001Δ*gdpd*_Xhp1_/TM7x compared with XH001/TM7x. (B) RNA-seq heatmap showing differential expression of genes involved in cell envelope and cell wall-associated pathways in XH001Δ*gdpd*_Xhp1_/TM7x compared with XH001/TM7x. (C) MIC analysis of WT XH001 and XH001Δ*gdpd*_Xhp1_ strains against vancomycin and teicoplanin. Statistical significance was determined by two-tailed unpaired student’s *t-*test; *P* < 0.05 (*), *P* < 0.01 (**). (D) Representative fluorescence microscopy images showing GFP-labeled TM7x associated with XH001 and XH001Δ *gdpd*_Xhp1_ host cells. Scale bar = 5 μm. (E) Quantification of the percentage of host cells associated with TM7x in XH001/TM7x and XH001Δ*gdpd*_Xhp1_/TM7x cocultures from nine independent fields of view. Statistical significance was determined using two-tailed unpaired Student’s *t*-test; *P* < 0.0001(****). (F) Relative rhamnose-to-glucose/mannose ratio of purified cell wall carbohydrates measured using a modified anthrone assay. Statistical significance was determined using two-tailed unpaired Student’s *t*-test; *P* < 0.01 (**). (G) RNA-seq heatmap showing reduced expression of type IV pilus-associated genes in XH001Δ*gdpd*_Xhp1_/TM7x compared with XH001/TM7x. Data are shown from three biological replicates. RNASeq data in (A), (B), and (G) are shown from three biological replicates. (H) Survival of XH001/TM7x and XH001Δ*gdpd*_Xhp1_/TM7x cocultures following 24 h incubation in nutrient-free PBS. Cell viability was determined by CFU enumeration and expressed as a percentage of viable cells remaining relative to the initial inoculum (T = 0 h). Data were log₁₀-transformed prior to analysis. Statistical significance was determined using Welch’s *t*-test; *P* < 0.01 (**). (I) TM7x abundance in XH001/TM7x and XH001Δ*gdpd*_Xhp1_/TM7x cocultures following 24 h incubation in nutrient-free PBS. Statistical significance was determined using two-way ANOVA with genotype and starvation time as factors, followed by Tukey’s multiple-comparisons test; *P* <0.0001 (****), ns = not significant.

Transcriptomic analysis also revealed that deletion of *gdpd*_Xhp1_ altered the expression of numerous genes encoding functions associated with cell envelope modeling, such as peptidases, cell-surface proteins, polysaccharide biosynthesis enzymes and peptidoglycan-associated factors (Fig. 4B). To determine whether these transcriptional changes corresponded to functional alterations of the envelope, we measured susceptibility to the cell-wall-targeting antibiotics vancomycin and teicoplanin. The XH001Δ*gdpd*_Xhp1_ mutant displayed significantly increased sensitivity to both antibiotics relative to WT XH001 (Fig. 4C), suggesting altered envelope integrity.

Given the importance of the cell envelope in mediating TM7x-host interactions^15^, we next tested whether these envelope alterations affected episymbiotic colonization by TM7x. Fluorescence microscopy of GFP-labelled TM7x revealed abundant epibionts associated with WT XH001, whereas substantially fewer TM7x cells were detected on the Δ*gdpd*_Xhp1_ mutant (Fig. 4D). Quantification confirmed a significant reduction in TM7x abundance in cocultures containing the mutant host (Fig. 4E), demonstrating that GDPDXhp1 promotes cell envelope remodeling and facilitate efficient TM7x colonization.

To identify envelope features associated with this colonization defect, we analyzed cell-wall carbohydrate composition in host cells grown in the absence of TM7x. Relative carbohydrate profiling revealed a significant decrease in the rhamnose-to-glucose/mannose ratio in the Δ*gdpd*_Xhp1_ mutant compared with WT XH001 (Fig. 4F), indicating substantial remodeling of rhamnose-rich cell-surface glycan(s). These alterations were accompanied by reduced TM7x colonization, suggesting that GDPD_Xhp1_-dependent envelope remodeling influences the host surface landscape encountered by the epibiont. Given the reported role of cell-surface glycans in supporting TM7x association^13^, the altered glycan composition, including the decreased rhamnose-to-glucose/mannose ratio in the Δ*gdpd*_Xhp1_ mutant may contribute to its impaired colonization phenotype.

We then asked whether host remodeling influenced transcriptional responses in TM7x. RNA-seq analysis revealed significant downregulation of multiple type IV pilus (T4P) genes, including *pilZ2, pilB2, pilW2, pilV2* and *pilA1*^43^, when TM7x was associated with the Δ*gdpd*_Xhp1_ mutant host (Fig. 4G). Since T4P is required for TM7x-host interactions^43^, these changes are consistent with the reduced colonization observed on the mutant host.

Finally, our previous work showed that LD accumulation enhances starvation tolerance in XH001^15^, we assessed the impact of GDPD_Xhp1_ on symbionts fitness during nutrient limitation. Overnight-grown WT XH001/TM7x and XH001Δ*gdpd*_Xhp1_/TM7x cocultures were washed and incubated in nutrient-free PBS, and viability was quantified at 0 and 24 h. Following 24 h of starvation, the Δ*gdpd*_Xhp1_ host exhibited a significantly greater loss of viability than the WT XH001, displaying an approximately 40-fold greater reduction in viability (Fig. 4H and Supplementary Fig. 5A). Consistent with this result, SYTOX Green staining revealed a substantially higher proportion of membrane-compromised Δ*gdpd*_Xhp1_ mutant cells than WT XH001 after 24 h of starvation (Supplementary Fig. 5B, C). Notably, TM7x abundance increased significantly in WT XH001/TM7x co-culture during starvation, indicating that the epibiont can persist and proliferate when associated with host bacteria, despite the absence of exogenous nutrients. In contrast, no significant increase in TM7x abundance was detected in XH001Δ*gdpd*_Xhp1_/TM7x cocultures (Fig. 4I and Supplementary Fig. 5B).

Together, these results identify GDPD_Xhp1_ as a key determinant of host envelope homeostasis and demonstrate that prophage-derived glycerophosphodiester metabolism influences cell-surface organization, TM7x colonization, and survival during nutrient stress.

### GDPD_Xhp1_ is required for productive Xhp1 infection

Upon induction, Xhp1 undergoes lytic replication, and generates phage particles capable of infecting prophage-free strain XH001ΔXhp1^31, 32^. Genes encoding putative GDPDs have been well-documented among phages infecting Gram-positive bacteria^19–24, 29, 30^. In bacteriophage Ld17, which infects *Lactobacillus delbrueckii*, the virion baseplate-associated enzyme GDPD_Ld17_ degrades the polyglycerol phosphate backbone of cell wall teichoic acid (CWTA), releasing G3P and promoting productive infection^30^. We first showed that deletion of *gdpd*_xhp1_ did not significantly affect prophage inducibility and assembly (Supplementary Fig. 6). To determine whether GDPD_Xhp1_ plays a role in Xhp1 infectivity, we examined its localization. Immunogold electron microscopy showed that GDPD_Xhp1_ is associated with the Xhp1 baseplate, whereas no labeling was observed in Xhp1Δ*gdpd* mutant phage particles (Fig. 5A). Despite normal phage assembly, deletion of *gdpd*_Xhp1_ abolished plaque formation on susceptible host bacterium XH001ΔXhp1, in contrast to WT Xhp1, which produced clear plaques indicative of productive infection (Fig. 5B). Together, these data indicate that GDPD_Xhp1_ is a baseplate-associated factor required for Xhp1 infectivity but not phage activation and assembly.

**Figure 5:**
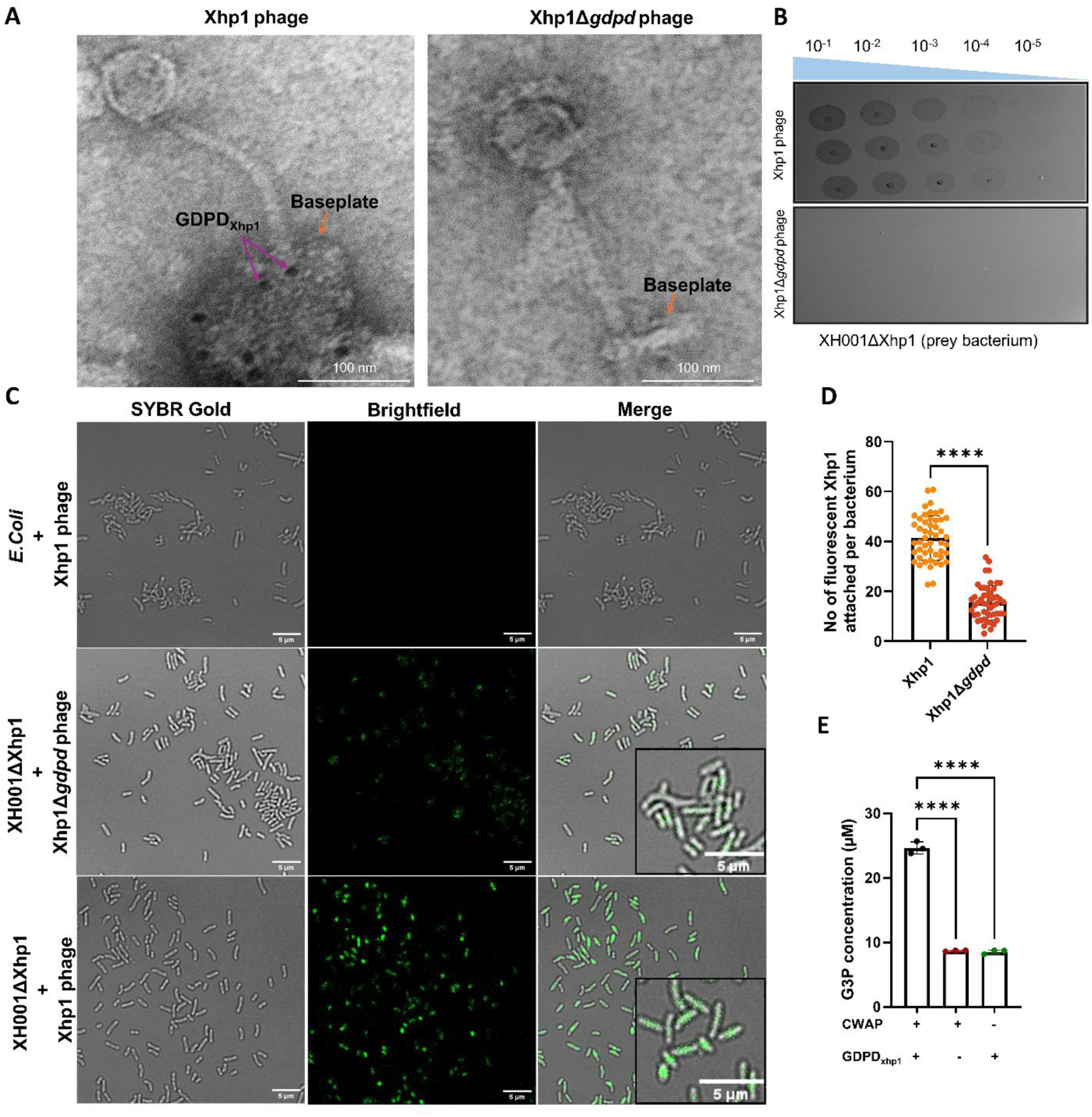
GDPD_Xhp1_ promotes phage infectivity through cell-wall polysaccharide remodeling and modulates host susceptibility. (A) Transmission electron micrographs showing immunogold-labeled GDPD_Xhp1_ localized to the baseplate of Xhp1 phage. Purple arrows indicate GDPD_Xhp1_, and orange arrows indicate the phage baseplate. (B) Plaque assay analysis showing impaired infectivity of Xhp1Δ*gdpd* phages. (C) Adsorption analysis of WT Xhp1 and Xhp1Δ*gdpd* phages. SYBR Gold-based visualization demonstrated reduced association of mutant phages with the prophage-free host strain XH001ΔXhp1 relative to WT Xhp1. (D) Quantification of phage attachment confirmed the reduced association of Xhp1Δ*gdpd* phages with XH001ΔXhp1 cells. Statistical significance was determined by Welch’s *t-*test; *P* < 0.0001 (****). (E) Biochemical characterization of GDPD_Xhp1_ activity. Purified GDPD_Xhp1_ hydrolyzed cell-wall-associated polysaccharides (CWAP) isolated from XH001 monocultures, releasing G3P from complex carbohydrate substrates. Statistical significance was determined using one-way ANOVA *P* < 0.0001 (****).

To determine whether the infectivity defect resulted from impaired host recognition, we examined phage adsorption by fluorescence microscopy using SYBR Gold-stained virions (Fig. 5C). No detectable phage attachment was observed on the non-host control *E. coli*. In contrast, WT Xhp1 bound efficiently to the prophage-free host strain XH001ΔXhp1, producing abundant surface-associated fluorescence. Although Xhp1Δ*gdpd* phage particles also associated with XH001ΔXhp1 cells, fluorescence signals were substantially reduced relative to WT Xhp1. Quantification confirmed a significant decrease in phage attachment, with WT Xhp1 averaging approximately 40 surface-associated particles per bacterium, whereas substantially fewer Xhp1Δ*gdpd,* around 15 particles per bacterium, were detected (Fig. 5D).

Successful phage infection often relies on enzymatic degradation or remodeling of cell-surface structures to increase access to host receptors and facilitate genome translocation across the cell envelope^44^. In *Actinomyces* and related Gram-positive bacteria, CWTAs are anionic phosphodiester-linked polyglycerol phosphate polymers integrated into the peptidoglycan matrix^45–47^, and genes required for CWTA biosynthesis are present in XH001^48^. Using cell-wall-associated polysaccharides (CWAP), a heterogeneous cell-envelope fraction isolated from XH001 as substrates, we found that recombinant GDPD_Xhp1_ released G3P (Fig. 5E), consistent with GDPD activity and resembling the activity reported for GDPD_Ld17_^30^. Together, these findings suggest that GDPD_Xhp1_ promotes productive infection by remodeling CWTA-containing cell-envelope structures, thereby enhancing phage-host interactions.

## Discussion

Prophages are increasingly recognized as important contributors to bacterial adaptation through the transfer of regulatory circuits, toxins, or defense systems that enhance host fitness under specific environmental conditions^4, 5, 49–51^. However, examples in which prophage genes become integrated into core metabolic processes remain comparatively rare. Here, we identify a prophage-derived glycerophosphodiester phosphodiesterase, GDPD_Xhp1_, that has been functionally domesticated by *S. odontolytica* XH001 and evolved as a central regulator of lipid metabolism, cell-envelope architecture, episymbiotic interaction, and phage fitness. Our findings reveal that a single viral gene can simultaneously influence the physiology of the bacterial host, the persistence of its episymbiont TM7x, and the reproductive success of the prophage from which it originated. This convergence of bacterial, episymbiotic, and viral fitness expands current concepts of prophage domestication and suggests that metabolic integration may represent an underappreciated outcome of long-term phage–host coevolution.

A central finding of this study is that GDPD_Xhp1_ is necessary for LD biogenesis in XH001. TM7x association strongly induced *gdpd*_Xhp1_ expression, and deletion of either *gdpd*_Xhp1_ or the entire prophage abolished TM7x-dependent LD accumulation (Fig. 1A-B and 2D-E). Lipidomic analyses further showed that GDPD_Xhp1_ promotes the reallocation of membrane glycerophospholipids into neutral storage lipids, particularly triacylglycerols and diacylglycerols (Fig. 2H), the major constituents of actinobacterial LD^36^. Notably, *gdpd*_Xhp1_ was expressed at substantially higher levels than the two chromosomally encoded GDPD homologues (Fig. 3A), arguing against functional redundancy and instead suggesting that XH001 has become reliant on a prophage-derived enzyme for maintaining lipid metabolic homeostasis. These findings provide a mechanistic explanation for the extensive lipid remodeling previously observed during TM7x episymbiosis^15^ and identify a specific prophage-encoded catalyst underlying this phenotype. More broadly, they demonstrate that prophage domestication can directly reshape central metabolic processes, extending the influence of prophages beyond canonical roles in defense, virulence, and horizontal gene transfer into the regulation of core bacterial physiology.

The metabolic consequences of GDPD_Xhp1_ activity appear to be linked to its role in generating G3P. G3P occupies a critical position at the intersection of phospholipid turnover, carbon metabolism, and respiratory electron transfer^17, 18^. Consistent with this role, disruption of *gdpd*_Xhp1_ reduced intracellular G3P abundance and was accompanied by extensive transcriptional reprogramming of respiratory pathways, including genes encoding NADH-quinone oxidoreductase and nitrate reductase complexes (Fig. 4A). Although our data do not establish a direct causal relationship between reduced G3P levels and the observed transcriptional changes, they support a model in which GDPD_Xhp1_ influences metabolic and redox homeostasis through regulation of G3P availability. In this context, TM7x-induced activation of GDPD_Xhp1_ may represent a strategy through which the host reallocates membrane-derived carbon to accommodate the increased energetic demands associated with episymbiosis.

Beyond its metabolic functions, GDPD_Xhp1_ emerged as an important determinant of membrane homeostasis. Deletion of *gdpd*_Xhp1_ altered membrane composition, increased membrane rigidity (Fig. 3C), and changed susceptibility to membrane and cell wall-targeting antibiotics in XH001 (Fig.3E). These findings reinforce growing evidence that bacterial membranes are highly adaptive structures whose physical properties are tightly linked to metabolic state^18, 52, 53^. Interestingly, loss of GDPD_Xhp1_ shifted phosphatidylcholine composition toward ether-linked species typically associated with increased membrane order (Fig. 3D). Such compensatory remodeling likely contributes to the elevated membrane rigidity observed in the mutant and may underlie the differential susceptibility to daptomycin antibiotics. Together, these results place a prophage-derived enzyme upstream of multiple aspects of membrane physiology and suggest that viral contributions to bacterial envelope homeostasis may be more extensive than currently appreciated.

Our results further reveal that GDPD_Xhp1_ influences organization of the bacterial cell surface. Loss of the enzyme altered the relative abundance of rhamnose-containing cell-wall carbohydrates, resulting in a decreased rhamnose-to-glucose/mannose ratio (Fig. 4F) and impaired TM7x colonization (Fig. 4D, E). These findings suggest that GDPD_Xhp1_-dependent metabolic remodeling extends beyond membrane lipid composition to influence the architecture of the cell envelope. Although the molecular mechanisms linking phospholipid turnover to rhamnose-containing cell-wall polysaccharide biosynthesis remain unresolved, envelope biogenesis is increasingly recognized as an integrated network in which membrane composition and cell-wall assembly are coordinately regulated^47^. We therefore propose that GDPD_Xhp1_-dependent remodeling alters the abundance, accessibility, or spatial organization of host surface determinants required for TM7x attachment. Consistent with this interpretation, TM7x associated with the Δ*gdpd*_Xhp1_ mutant exhibited reduced abundance and downregulated genes involved in type IV pilus biogenesis (Fig. 4G), structures known to mediate host interaction^43^. These findings suggest that GDPD_Xhp1_-dependent metabolic remodeling can influence TM7x colonization and may alter the physical interface underlying bacterial episymbiosis.

The starvation phenotype of XH001Δ*gdpd*_Xhp1_/TM7x co-culture further underscores the role of GDPD_Xhp1_ in maintaining the XH001–TM7x association under nutrient limitation (Fig.4H, I). Previous work showed that LD accumulation enhances starvation tolerance in XH001^15^, suggesting that the reduced survival of XH001 Δ*gdpd*_Xhp1_ is linked to its defect in LD biogenesis. Notably, TM7x abundance increased in WT cocultures despite the absence of exogenous nutrients for 24 h, indicating that the epibiont can persist and proliferate while associated with a viable host. In contrast, TM7x failed to expand in XH001Δ*gdpd*_Xhp1_/TM7x cocultures, suggesting that host metabolic fitness is a key determinant of episymbiont persistence under starvation. Together, these findings support a model in which GDPD_Xhp1_-mediated lipid remodeling promotes host resilience during nutrient stress and thereby stabilizes the XH001–TM7x interaction. Whether TM7x actively induces this metabolic state or instead benefits from a host adaptation remains an important question for future investigation.

Remarkably, GDPD_Xhp1_ also retained an essential function in the phage life cycle (Fig. 5). Recombinant GDPD_Xhp1_ hydrolyzed cell-wall-associated substrates and released G3P, consistent with degradation of phosphodiester-linked polymers within the Gram-positive cell envelope^30^. This activity parallels that of virion-associated GDPDs reported in other bacteriophages^30^ and support a model in which phage-associated GDPD_Xhp1_ remodels host cell-envelope structures to enhance phage-host interactions and promote productive infection. The preservation of this ancestral phage function alongside the acquisition of host-beneficial metabolic activities highlights a striking example of prophage domestication. Rather than becoming dispensable following lysogenization, GDPD_Xhp1_ appears to have evolved into a multifunctional enzyme that benefits both the bacterial host and the prophage, coupling host metabolism with phage fitness.

In summary, we demonstrate that a prophage-derived GDPD_Xhp1_ has been evolved as a central regulator of bacterial physiology, linking lipid metabolism, cell envelope architecture, episymbiotic interaction, and phage reproduction (Fig. 6). GDPD_Xhp1_ represents an unusual mode of prophage domestication in which a retained viral function simultaneously enhances host adaptation, supports episymbiont growth, and contributes to phage fitness. Our study raises several important questions for future investigation. Although TM7x episymbiosis induces *gdpd*_Xhp1_ expression, the underlying regulatory mechanisms remain unknown. Likewise, it is unclear how the prophage-derived *gdpd*_Xhp1_ became functionally integrated into host metabolism and emerged as the predominant GDPD-encoding gene despite the presence of two chromosomal GDPD homologues in XH001. Future studies addressing these questions will clarify how symbiotic interactions drive prophage domestication and illuminate the evolutionary processes through which prophage-encoded functions are integrated into core bacterial physiology. Nevertheless, our findings expand the conceptual framework of prophage domestication beyond conventional accessory traits and reveal how viral genes can become deeply integrated into metabolic and ecological networks of bacterial cells. Given the widespread distribution of prophage-encoded metabolic enzymes across bacterial genomes^54, 55^, similar examples of metabolic domestication may be more common than currently appreciated. Collectively, our results highlight prophages as important drivers of metabolic innovation that can shape host physiology, microbial interactions, and evolutionary trajectories.

**Figure 6.**
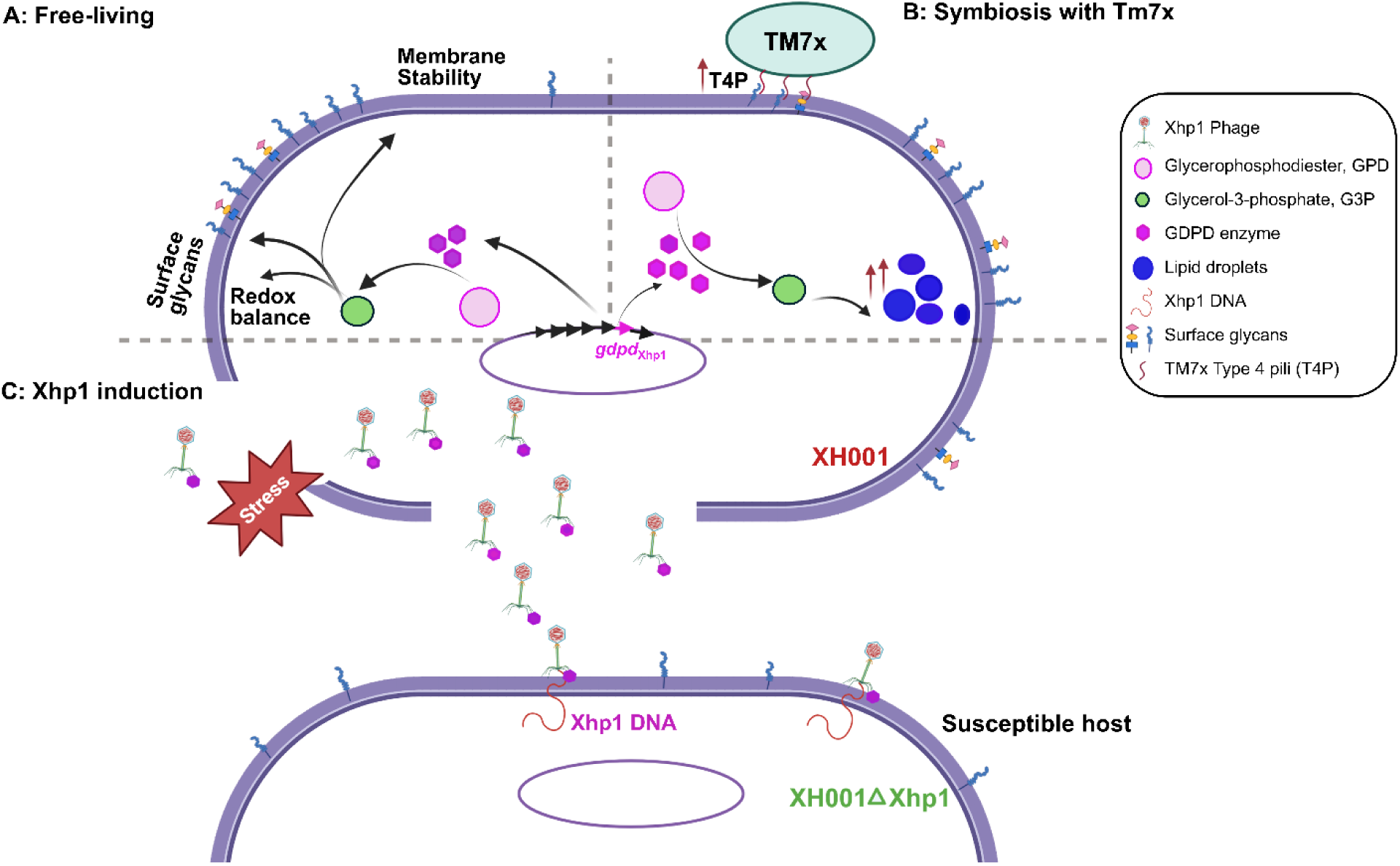
Proposed model for the multifunctional roles of prophage-derived GDPD_Xhp1_ in host physiology, TM7x episymbiosis and Xhp1 infection. (A) In XH001, GDPD_Xhp1_ catalyzes glycerophosphodiester phosphodiesterases (GPD) turnover to generate glycerol-3-phosphate (G3P), a central metabolic intermediate linking phospholipid metabolism to cellular energy production, thereby supporting membrane homeostasis, redox balance, and cell-envelope organization. (B) During episymbiosis with TM7x, GDPD_Xhp1_ promotes lipid droplet biogenesis in XH001, facilitates TM7x colonization and enhances expression of type IV pilus (T4P)-associated genes in TM7x. (C) Following induction of the Xhp1 prophage, GDPD_Xhp1_ retains its ancestral phage function by remodeling cell wall teichoic acid (CWTA)-containing cell-envelope structures, thereby enhancing phage-host interactions and promoting productive infection. Together, these findings support a model in which GDPD_Xhp1_ functions as a central metabolic node coordinating bacterial physiology, episymbiotic interactions, and phage fitness.

## Materials and methods

### Bacterial strains, prophages, and growth conditions

*N. lyticus* TM7x (type strain TM7x HMT-952) and its host bacterium *S. odontolytica* XH001 were originally co-isolated from the human oral cavity. Cultures containing both organisms are referred to as XH001/TM7x co-cultures. The GFP-labeled TM7x strain, *N. lyticus* NS1::sfGFP, was used for microscopic visualization and fluorescence-based assays^56^. All cultures were grown in Brain Heart Infusion (BHI; BD Biosciences) medium, with or without 5% defibrinated sheep blood, at 37 °C under micro-oxic conditions (2% O₂, 5% CO₂, 93% N₂) using a Whitley A35 workstation (Don Whitley Scientific)^43, 56^. For experiments requiring anoxic conditions, cultures were maintained in an anaerobic chamber (Coy Laboratories, Grass Lake, MI, USA) supplied with a gas mixture of 5% H₂, 10% CO₂, and 85% N₂. Complemented strains were cultured in BHI supplemented with kanamycin (400 μg/mL).

### Construction and validation of *gdpd*_Xhp1_ knockout mutants

*S. odontolytica* XH001 was made electrocompetent and genetically manipulated as previously described^57–59^. To generate an allelic exchange construct targeting the *gdpd*_Xhp1_ locus (APY09_RS09270), approximately 500-bp regions flanking the target gene were PCR-amplified. The upstream and downstream homology arms were assembled with the thiamphenicol-resistance cassette (*catP*) derived from pCWU6^60^ and cloned into the shuttle plasmid pCWU3^61^. For electroporation, XH001 cultures were grown to the mid-logarithmic phase and treated with 3.75% (w/v) glycine to increase cell wall permeability. Cells were subsequently harvested, washed once with sterile water and twice with 10% glycerol, and finally resuspended in 10% glycerol. The resulting electrocompetent cells were mixed with 1–2 µg of linearized DNA and electroporated at 2.5 kV, 25 µF, and 400 Ω. Following electroporation, cells were plated on BHI agar supplemented with thiamphenicol (10 µg mL⁻¹) and incubated for 24 h. Putative transformants were subsequently replica-plated onto BHI agar containing either thiamphenicol (10 µg mL⁻¹) or kanamycin (400 µg mL⁻¹). Colonies that retained thiamphenicol resistance but were unable to grow in the presence of kanamycin were considered candidate double-crossover mutants and were selected for validation.

### Construction of *gdpd*_Xhp1_ complemented strains

Genetic complementation was performed following a previously described strategy^62^. For genetic complementation, a DNA fragment encompassing the *gdpd*_Xhp1_ coding sequence along with 100 bp upstream of the predicted start codon, including the native ribosome-binding site, was PCR-amplified (Supplementary Table 1) from XH001 genomic DNA using Q5 High-Fidelity DNA Polymerase (NEB, Ipswich, MA). The amplified fragment was inserted into XhoI-digested pJRD215 using the NEBuilder HiFi DNA Assembly Cloning Kit (NEB, Ipswich, MA). The resulting plasmid construct was introduced into XH001 and XH001Δ*gdpd*_Xhp1_ strains. Transformants were selected on kanamycin-supplemented BHI agar, validated by PCR, and analyzed for *gdpd*_Xhp1_ expression by quantitative real time PCR (Supplementary Table 1).

### Phage induction, purification, and titer determination

Prophages were induced from mid-log-phase *S. odontolytica* cultures (OD₆₀₀ ≈ 0.5) using mitomycin C (0.01 µg mL⁻¹) and incubated for 6–24 h. Following induction, cultures were clarified by centrifugation (8,000 × g for 10 min) and the supernatant was filtered through 0.22 µm membranes. Phage particles were precipitated by adding NaCl (1 M final concentration) and polyethylene glycol 8000 (10% w/v) and incubating overnight at 4 °C. The precipitate was recovered by centrifugation (10,000 × g for 30 min), resuspended in SM buffer (50 mM Tris-HCl [pH 7.5], 100 mM NaCl, 8 mM MgSO₄), and further purified by CsCl density-gradient ultracentrifugation (100,000 × g for 2 h at 4 °C). Phage bands were collected, dialyzed against SM buffer, and stored at 4 °C^63^. Phage titers were determined using the double-layer agar assay^64^. Briefly, a lawn of host cells (XH001ΔXhp1) was prepared in 0.3% molten soft agarose and overlaid onto BHI agar plates. Tenfold serial dilutions of phage lysates were prepared in SM buffer. Aliquots of each phage dilution were then spotted directly onto the prepared host-cell lawn and allowed to absorb for 10 min. The plates were incubated for 24 h, after which plaques were enumerated. Plaques were counted and titers were calculated as plaque-forming units per milliliter (PFU mL⁻¹)^65^.

### Expression and purification of GDPD_Xhp1_ protein

The full-length *gdpd*_Xhp1_ coding sequence was cloned (Supplementary Table 1) into the pET28a expression vector to generate an N-terminal His-tagged fusion protein. The construct was sequence-verified and transformed into *E. coli* BL21(DE3) for protein overexpression. Cultures were grown under standard induction conditions with 0.5 mM IPTG, harvested by centrifugation at 5,000 × g for 10 min at 4 °C, and cell pellets were resuspended in lysis buffer consisting of 50 mM NaH₂PO₄, 300 mM NaCl, and 20 mM imidazole (pH 8.0). Cells were lysed by sonication on ice at 50% amplitude using 10-s pulses for a total sonication time of 60 s per sample. Lysates were clarified by centrifugation at 12,000 × g for 10 min at 4 °C, and the soluble fraction was retained for purification. His-tagged GDPD_Xhp1_ was purified by immobilized metal affinity chromatography (IMAC) using an ÄKTA fast protein liquid chromatography (FPLC) system pre-equilibrated with IMAC binding buffer (50 mM NaH₂PO₄, 300 mM NaCl, 20 mM imidazole, pH 8.0). Clarified lysates were loaded onto a Ni²⁺-charged affinity column, washed extensively with binding buffer to remove nonspecifically bound proteins, and eluted with buffer containing 300 mM imidazole. Elution fractions were monitored by absorbance at 280 nm, collected, and analyzed by SDS-PAGE. Fractions containing purified GDPD_Xhp1_ were pooled, dialyzed, and concentrated using a 10-kDa molecular-weight-cutoff centrifugal concentrator (Sigma-Aldrich, Saint Louis, MO).

### Fluorescence staining and confocal microscopy

Neutral lipid accumulation was visualized in live cells using LipidSpot staining. Samples were incubated with LipidSpot^TM^ dye (Biotium) at the manufacturer-recommended 1× working concentration for 3 h. Stained cells were immediately imaged without fixation. Fluorescence excitation was performed at 488 nm, and emission was collected between 500 and 550 nm. Membrane fluidity was assessed using the membrane-sensitive dye Laurdan (6-dodecanoyl-2-dimethylaminonaphthalene; Thermo Fisher Scientific) under super-resolution Airyscan imaging. Laurdan was prepared in dimethylformamide (DMF; Sigma-Aldrich), maintaining a final DMF concentration of 1% (v/v) in the staining medium. Cells were grown in BHI supplemented with daptomycin and 1.25 mM CaCl₂, washed twice with pre-warmed PBS, and incubated with Laurdan for 30 min. Following staining, cells were washed twice with 1× PBS before imaging.Laurdan was excited at 405 nm, and emission intensities were collected at 460 ± 5 nm (for liquid ordered phase) and 500 ± 5 nm (for liquid disordered phase), respectively. Generalized polarization (GP) values were calculated using the equation: GP = (I₄₆₀ − I₅₀₀)/(I ₄₆₀ + I₅₀₀). I_460_ and I_500_ were the fluorescence intensity from the abovementioned two phases, respectively.

For visualization of isolated *gdpd*_Xhp1_ phage particles and phage–host association, purified phages (Xhp1 and Xh001Δ*gdpd*) preparations were stained with SYBR Gold nucleic acid dye according to the manufacturer’s recommended working concentration. Samples were protected from light during staining. SYBR Gold-labeled phage particles were subsequently mixed with XH001ΔXhp1 host cells and immediately visualized by confocal microscopy without fixation. SYBR Gold fluorescence was excited at 488 nm, and emission was collected between 500 and 550 nm.

For visualization of cell damage and loss of membrane integrity, XH001/TM7x and Δ*gdpd*_Xhp1_/TM7x cocultures were stained with SYTOX Green nucleic acid stain according to the manufacturer’s recommended working concentration. Briefly, cells were collected at the indicated time points, washed with PBS, and resuspended in PBS. SYTOX Green fluorescence was subsequently visualized by confocal microscopy without fixation. The fluorophore was excited at 488 nm, and emission was collected between 500 and 550 nm. SYTOX Green-positive cells, which indicate compromised plasma membrane integrity and loss of viability, were used as a measure of cell damage/death.

### Minimum inhibitory concentration (MIC) assay

For minimum inhibitory concentration (MIC) determination, bacterial cultures were adjusted to an initial optical density (OD₆₀₀) of 0.1 and dispensed into sterile 96-well microplates (Thermo Fisher Scientific, MA). Antibiotics were serially diluted in the wells to final concentrations ranging from 1024 to 1 µg/ mL. Plates were incubated overnight at 37 °C under micro-oxic conditions. Following incubation, bacterial growth was quantified by measuring OD₆₀₀ using an iMark™ microplate absorbance reader (Bio-Rad). The MIC was defined as the lowest antibiotic concentration at which no detectable bacterial growth was observed.

### Quantitative Real-Time PCR (qRT-PCR)

Bacterial cultures were grown in BHI medium at 37 °C to mid-exponential phase (OD₆₀₀ ≈ 0.3– 0.4). Total RNA was isolated from 2 mL of culture using the MasterPure Complete DNA and RNA Purification Kit (EPICENTRE; Cat. Nos. MC85200 and MC89010) according to the manufacturer’s instructions. Purified RNA was reverse-transcribed into complementary DNA (cDNA) using the HiFiScript cDNA Synthesis Kit (CoWin BioSciences, Cambridge, MA, USA). Quantitative PCR reactions were performed in triplicate using 10 ng of cDNA template, 10 µM of each gene-specific primer, and PowerUp™ SYBR™ Green Master Mix (Applied Biosystems). Amplification was carried out on a QuantStudio™ 3 Real-Time PCR System (Thermo Fisher Scientific, Waltham, MA, USA) under the following conditions: an initial denaturation at 95 °C for 1 min, followed by 40 cycles of 95 °C for 30 s and 60 °C for 30 s. Transcript levels were normalized to 16S rRNA, and relative gene expression was calculated using the 2^⁻ΔΔCT^ method. Primer sequences used in this study are listed in supplementary Table 1.

### Raman spectroscopy

Raman spectroscopy was used to characterize the cellular chemical composition following established approaches commonly used in microbiological studies^66^. Cells were fixed in 10% neutral buffered formalin (Sigma-Aldrich) for 60 min at room temperature, washed three times with sterile water, and air-dried onto aluminum-coated Raman substrates. Raman spectra were acquired using an HR Evolution confocal Raman microscope (Horiba Jobin-Yvon, France) equipped with a 532-nm Nd laser. The laser power at the sample was adjusted to 8 mW using 25% neutral density filters. A 100× objective (NA = 0.6) was used to focus the laser onto cell aggregates, generating an approximate spot size of 1 μm². Raman scattering was detected using a charge-coupled device (CCD) detector maintained at −70°C. The spectra were acquired in the range of 300–1800 cm^−1^. A 600 grooves/mm diffraction grating was used, providing a spectral resolution of approximately 1.5 cm⁻¹. Each spectrum was acquired with a 10-s integration time and two accumulations. Raw spectra were processed using LabSpec 6, including cosmic-ray removal, baseline correction, and vector normalization. Processed spectra were subsequently analyzed in RStudio (2024.09.1) using custom R scripts.

### Transmission Electron Microscopy

To visualize phage particles and examine phage–host interactions, samples from phage-free monocultures exposed to purified phage were processed for TEM. Cell pellets were fixed in 2.5% glutaraldehyde, 1.25% paraformaldehyde, and 0.03% picric acid in 0.1 M sodium cacodylate buffer (pH 7.4) for at least 2 h at room temperature. Following fixation, samples were washed with 0.1 M sodium cacodylate buffer and post-fixed in 1% osmium tetroxide containing 1.5% potassium ferrocyanide for 1 h. Samples were then washed twice with distilled water, rinsed once with maleate buffer, and stained with 1% uranyl acetate in maleate buffer for 1 h. After two additional washes with distilled water, samples were dehydrated through a graded ethanol series (50%, 70%, 90%, and twice in 100%; 10 min each), followed by incubation in propylene oxide for 1 h. Samples were infiltrated overnight with a 1:1 mixture of propylene oxide and Spurr’s low-viscosity resin (Electron Microscopy Sciences, Hatfield, PA, USA), embedded in fresh Spurr’s resin, and polymerized at 60°C for 48 h. Ultrathin sections (∼60 nm) were cut using a Reichert Ultracut-S ultramicrotome, collected on copper grids, and stained with lead citrate. Sections were examined using a Tecnai G2 Spirit BioTWIN transmission electron microscope (FEI), and digital images were acquired with an AMT NanoSprint 43-MKII camera.

### Immunogold TEM

The Xhp1-encoded glycerophosphodiester phosphodiesterase (GDPD_Xhp1_), a predicted baseplate-associated protein, was cloned, expressed, and purified as described above. The purified recombinant GDPD_Xhp1_ protein was used to generate a rabbit polyclonal antibody through Pacific Immunology Corp. (Pacific Immunology, CA). To assess the localization of GDPD_Xhp1_ within the Xhp1 baseplate, purified Xhp1 phages were subjected to immunogold labeling and examined by TEM. Briefly, samples were incubated with the rabbit polyclonal anti-GDPD_Xhp1_ primary antibody, followed by a 10-nm gold-conjugated goat anti-rabbit IgG secondary antibody. Following immunogold labeling and staining, the samples were examined by TEM to determine the localization of GDPD_Xhp1_ on the Xhp1 phage particles.

### Starvation assay

XH001/TM7x and Δ*gdpd*_Xhp1_/TM7x cocultures were initially cultured in a microaerophilic chamber for two passages. Following the second passage, cells were harvested, washed twice with sterile 1× PBS, and resuspended in PBS within the microaerophilic chamber to initiate nutrient starvation. Samples were collected immediately after resuspension (0 h) and following 24 h of incubation in PBS. At each time point, samples were analyzed for viable cell counts by colony-forming unit (CFU) assay, and for cell membrane integrity by SYTOX Green staining.

### Enzyme-coupled spectrophotometric assay

GDPD_Xhp1_ enzymatic activity was quantified by measuring G3P production using a two-step coupled spectrophotometric assay^30^. The 100 μL reaction mixture contained 50 mM MOPS (pH 7.5), 20 mM CaCl_2_, purified recombinant GDPD_Xhp1_, and substrate. Purified carbohydrates and cell wall-associated carbohydrate extracts were used at a final concentration of 2.5 g/L, whereas *L*-α-glycerophosphorylcholine (Sigma-Aldrich) was used at 1 mM. Reactions were incubated at 30 °C for 30 min and terminated by heating at 95 °C for 10 min before cooling to room temperature. G3P production was quantified using a Glycerol-3-Phosphate Assay Kit (MAK546, Sigma-Aldrich) according to the manufacturer’s instructions. Intracellular G3P levels were also measured using the same assay kit following the manufacturer’s protocol.

### Untargeted Lipidomics

Bacterial cultures were grown to an OD_600_ of 0.5, harvested by centrifugation and stored at −80°C until analysis. Lipids were extracted using methanol:acetonitrile:water (2:2:1, v/v/v), followed by protein normalization using a bicinchoninic acid (BCA) assay. Dried lipid extracts were reconstituted in acetonitrile:water (2:1, v/v) and analyzed by reversed-phase liquid chromatography–mass spectrometry (RPLC–MS) using a Waters Acquity Premier HSS T3 column (2.1 × 100 mm). Chromatographic separation was performed with water/acetonitrile/isopropanol mobile phases containing 10 mM ammonium formate at a flow rate of 400 μL min⁻¹ using a linear gradient. Mass spectra were acquired over an *m/z* range of 100– 1700 in both positive and negative ionization modes with data-dependent MS/MS acquisition. Raw data were processed using in-house software for peak detection, feature alignment, background filtering, and annotation. Metabolites were identified by comparison with the RefMet, LIPID MAPS, and HMDB databases using accurate mass, isotope pattern, retention time, and MS/MS fragmentation spectra, and annotated according to the Metabolomics Standards Initiative (MSI) guidelines. Features with a quality control coefficient of variation >25% were excluded, missing values were imputed using half of the minimum detected intensity, and metabolite intensities were log₂-transformed prior to statistical analysis. Differential metabolites were identified using Welch’s one-way ANOVA, and pathway enrichment analysis was performed using gene set enrichment analysis (GSEA) with Reactome pathways and LIPID MAPS lipid class annotations.

### RNA sequencing and data analysis

Total RNA was extracted using the MasterPure Complete DNA and RNA Purification Kit (EPICENTRE; Cat. Nos. MC85200 and MC89010) as described previously. RNA quality was assessed using a NanoDrop spectrophotometer (Thermo Fisher Scientific) and an Agilent 5400 system (Agilent Technologies, Santa Clara, CA, USA). Ribosomal RNA-depleted strand-specific RNA libraries were prepared using the NEBNext Ribo-Zero Magnetic Kit, generating libraries with 250–300 bp inserts. Paired-end sequencing was performed on the NovaSeq PE150 platform (Illumina, San Diego, CA, USA) at Novogene (Sacramento, CA, USA), producing 150 bp paired-end reads with approximately 2 Gb of sequencing output per sample.

Raw sequencing reads were mapped to a combined reference genome containing XH001 and TM7x sequences using a Bowtie2-based pipeline optimized for paired-end reads. Differential gene expression analysis was performed using the DESeq2 package in RStudio as previously described ^67^. All reported data are representative of three biological replicates.

### Cell wall isolation and modified anthrone assay

Bacterial cell wall purification was performed based on a previously described peptidoglycan isolation protocol^68^ with modifications to preserve cell wall-associated glycans. Briefly, 400 mL cultures of each strain were grown overnight. Cells were harvested, washed, frozen, and subsequently added dropwise into boiling PBS containing 8% (w/v) SDS for 30 min to lyse cells and isolate insoluble cell wall sacculi. The isolated sacculi were washed three times and sequentially treated with DNase (10 μg/mL), RNase (50 μg/mL), and Pronase (200 μg/mL) at 37 °C for 90 min each. Following enzymatic digestion, sacculi were frozen, resuspended in 1% PBS, and boiled for 30 min, followed by two washes with water, one wash with 100% acetone, and dehydration.

The modified anthrone assay was adapted from a previously reported method for analysis of cell wall polysaccharides in *Streptococcus mutans* ^69^. Briefly, anthrone reagent was prepared at 0.2% (w/v) solution in concentrated sulfuric acid. For each reaction, 1 mL of anthrone reagent was added to 200 μL of either monosaccharide standards (0.1% and 0.01% w/v) or purified sacculi suspensions (2 mg and 0.2 mg). Samples were incubated at 98°C for 10 min, cooled, and analyzed spectrophotometrically. Based on the absorption profiles of 6-deoxyhexoses (rhamnose and fucose) and non-6-deoxyhexoses (glucose and mannose), absorbance was measured at 580 nm (total carbohydrate signal) and 690 nm (non-6-deoxyhexose carbohydrate signal). The relative rhamnose-to-glucose/mannose ratio was calculated as: (OD_580_-OD_690_)/OD_690_.

### Extraction of cell wall associated carbohydrates

Cell wall-associated carbohydrates were extracted as previously described^30^. Briefly, a 400 mL culture of fresh medium was inoculated with 1% overnight bacterial culture and incubated for approximately 20 h at the appropriate growth temperature. Cells were harvested by centrifugation (8,700 × g, 15 min) and washed three times with 20 mM ammonium acetate buffer (NH₄C₂H₃O₂/C_2_H_4_O_2_, pH 4.7). The cell pellet was resuspended in the same buffer containing 4% SDS and boiled with vigorous stirring for 1.5 h. After cooling to room temperature, cells were collected by centrifugation (8,700 × g, 15 min) and washed three to five times with 20 mM ammonium acetate buffer (pH 4.7) until no SDS-mediated foaming was observed. The washed pellet was resuspended in 50 mL of 5% trichloroacetic acid containing acid-washed glass beads (Sigma-Aldrich) and stirred at 4°C for 48 h. Following centrifugation, the supernatant was dialyzed against distilled water at 4°C and lyophilized. The dried material was resuspended in 5 mL of water and centrifuged at 25,700 × g for 20 min to remove residual insoluble material. The clarified supernatant was collected, lyophilized, and stored at room temperature.

### Statistical analysis

Statistical analyses were conducted using GraphPad Prism (version 10.3.1). The choice of statistical test depended on the experimental design and dataset, with unpaired two-tailed *t*-tests, Welch’s *t*-test and one/two-way analysis of variance (ANOVA) applied as appropriate, followed by Tukey’s post hoc multiple-comparison test. Statistical significance was assigned based on the following thresholds: *P* < 0.05 (*), *P* < 0.01 (**), *P* < 0.001 (***), and *P* < 0.0001 (****).

## Acknowledgements

Research in this publication was supported by the National Institute of Dental and Craniofacial Research of the NIH under awards 1R01DE023810 (to X.H., J.S.M., B.B), R01DE030943 (to X.H.), 1R01DE031274 (to B.B.), and K99/R00 DE033794 (to P.-T.D). Microscopy was performed at the ADA Forsyth Institute Advanced Microscopy Core Facility (RRID:SCR_021121). Electron microscopy imaging was conducted at the Harvard Medical School electron microscopy facility. Lipidomics was performed and analyzed by Panome Bio (St. Louis, MO). Polyclonal antibody production was performed by Pacific Immunology (Ramona, CA). The content is solely the responsibility of the authors and does not necessarily represent the official views of the NIH.

## Authors’ contributions

Conceptualization was carried out by NN, PTD and XH. Methodology was developed by NN, AK, ASG, PTD and XH. NN and AK performed experiments, data acquisition, image acquisition, and data analysis. PTD assisted with Raman spectroscopy acquisition and data analysis. JSM, BB, and XH provided scientific input and contributed to the interpretation of the data. NN wrote the original draft. All authors discussed the results, reviewed, revised and approved of the final manuscript.

---

**Supplementary Figure 1:**
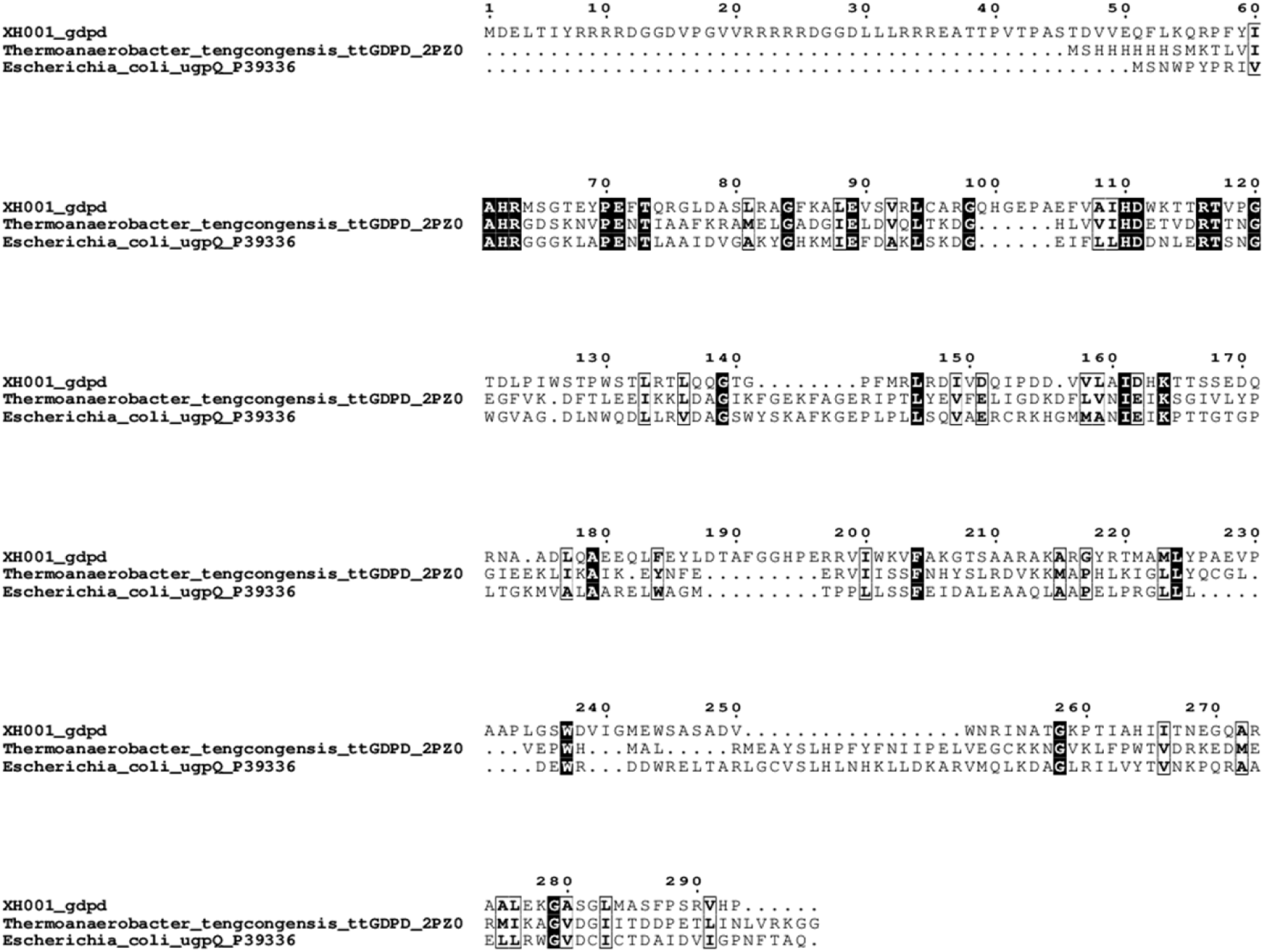
Multiple sequence alignment of GDPD_Xhp1_ enzymes. Multiple sequence alignment (MSA) of the prophage-encoded glycerophosphodiesterase GDPD_Xhp1_ with experimentally characterized homologues from *T. tengcongensis* (ttGDPD, PDB ID: 2PZ0) and *E. coli* (ugpQ, UniProt ID: P39336). The alignment was generated using Clustal Omega and formatted using ESPript 3.0. Identical residues across all three sequences are highlighted in black boxes with white text. Conserved residues with similar physicochemical properties are enclosed in thin black outlines. Dots (.) represent alignment gaps introduced to maximize sequence homology. Numbering at the top corresponds to the total alignment position.

**Supplementary Figure 2:**
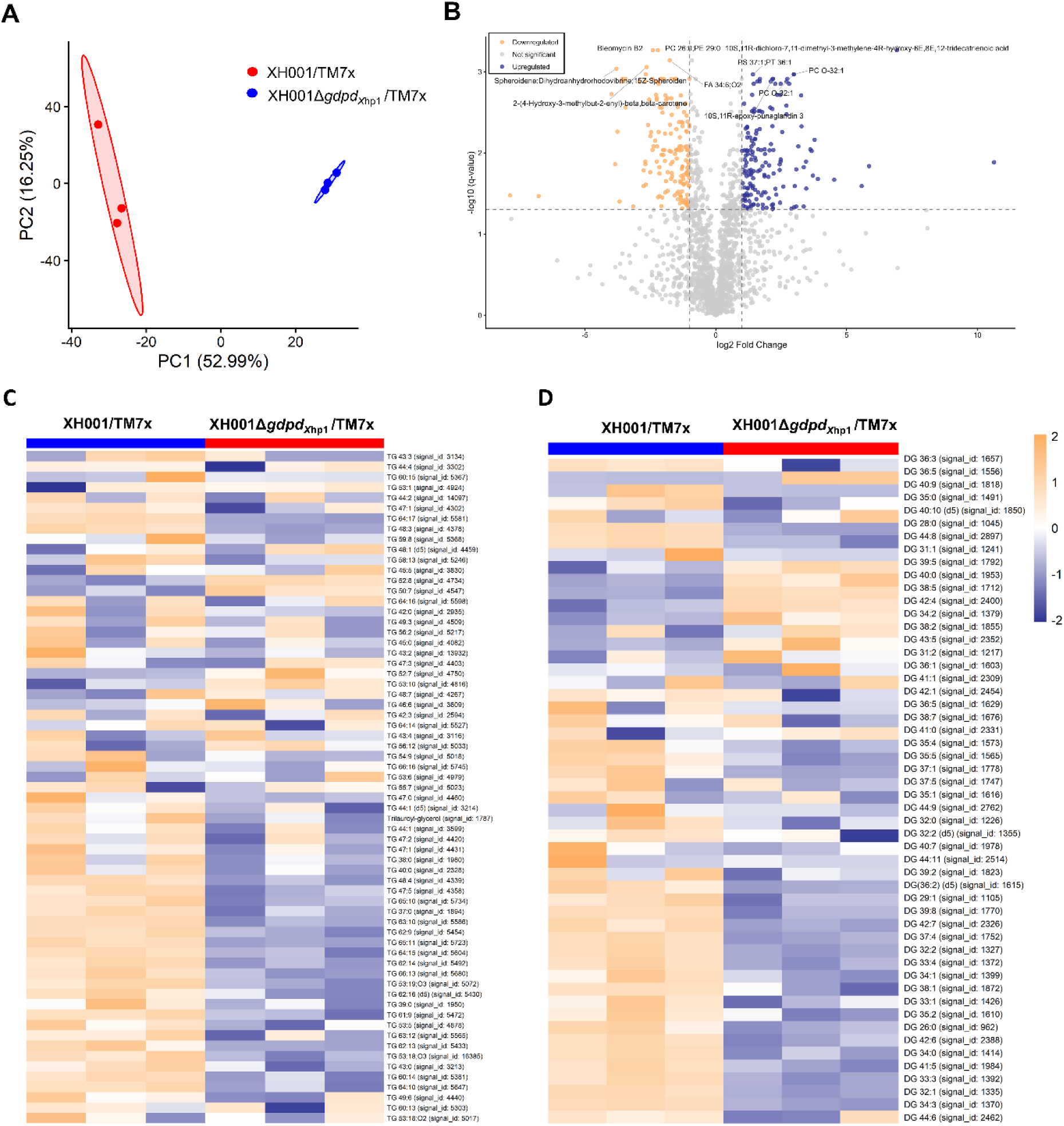
Global lipidomic profiling reveals extensive remodeling of neutral lipid metabolism in XH001/TM7x coculture. (A) PCA of lipidomic profiles demonstrating clear separation between XH001/TM7x and XH001Δ*gdpd*_Xhp1_/TM7x cocultures, indicating distinct lipid compositions. Ellipses represent the 95% confidence regions for each group. (B) Volcano plot showing differential lipid metabolite abundance in XH001Δ *gdpd*_Xhp1_/TM7x relative to XH001/TM7x. Log₂ fold changes are plotted against adjusted *P* values (q values). Significantly decreased and increased metabolites are highlighted in orange and blue, respectively. (C-D) Heatmaps of triacylglycerols (GL0301) (C) and diacylglycerols (GL0201) (D). Each row represents an individual lipid species, and each column represents a biological replicate. Color intensity indicates the relative abundance of each lipid across XH001/TM7x and XH001Δ*gdpd*_Xhp1_/TM7x cocultures. Loss of *gdpd*_Xhp1_ altered neutral lipid metabolism, resulting in changes in triacylglycerol and diacylglycerol profiles associated with LD regulation and cellular lipid homeostasis.

**Supplementary Figure 3:**
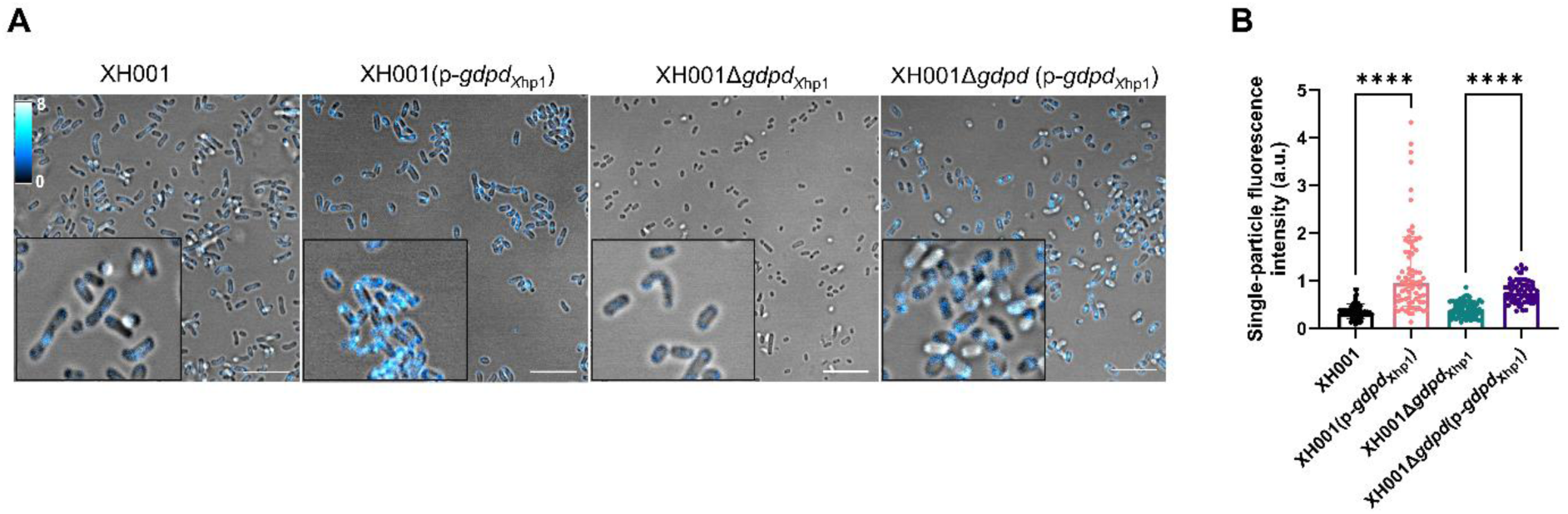
Complementation with *gdpd*_Xhp1_ rescues LDs defects in the Δ*gdpd*_Xhp1_ mutant. (A) Representative confocal laser scanning microscopy images of LipidSpot-stained lipid droplets in WT XH001, XH001Δ*gdpd*_Xhp1_, XH001(p-*gdpd*_Xhp1_), and XH001Δ*gdpd*(P-*gdpd*_Xhp1_) strains. Scale bar = 5 μm. (B) Quantification of LipidSpot fluorescence intensity in the indicated strains. Constitutive expression of *gdpd*_Xhp1_ restored LD accumulation in both complemented strains, demonstrating that GDPD_Xhp1_ is sufficient to drive LD formation independently of TM7x. Statistical significance was determined by one-way ANOVA with multiple-comparisons testing; ****, *P* < 0.0001.

**Supplementary Figure 4:**
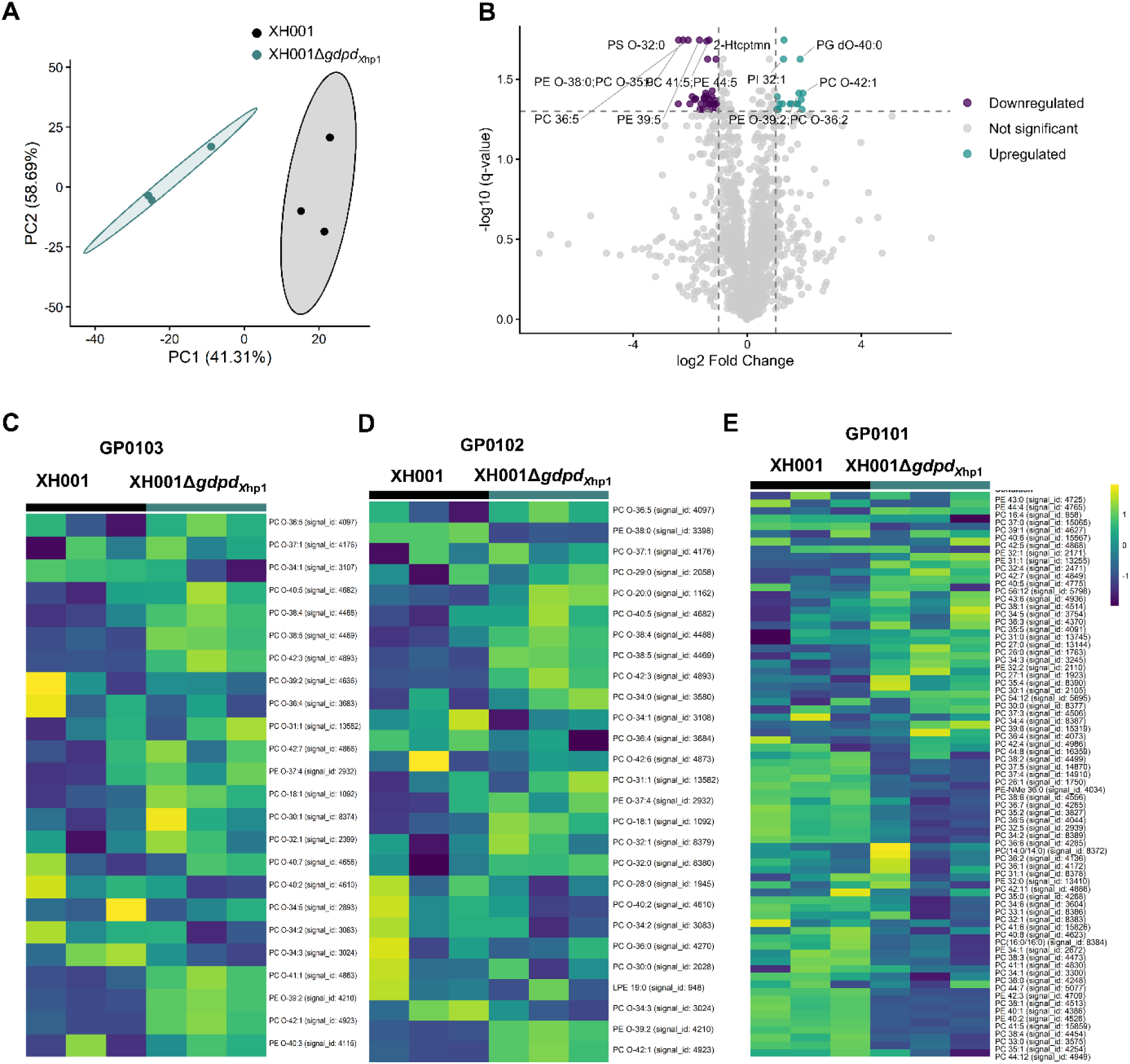
Global lipidomic profiling reveals phosphatidylcholine remodeling in XH001Δ*gdpd*_Xhp1_ monoculture. (A) PCA of lipidomic profiles demonstrating clear separation between XH001 and XH001Δ*gdpd*_Xhp1_ monocultures, indicating distinct lipid compositions following deletion of *gdpd*_Xhp1_. Each point represents an individual biological replicate, and ellipses indicate the 95% confidence regions of each group. (B) Volcano plot showing differential lipid metabolite abundance in XH001Δ *gdpd*_Xhp1_ relative to XH001. Log₂ fold changes are plotted against adjusted *P* values (q values). Significantly decreased and increased metabolites are highlighted in purple and yellow, respectively. (C-E) Heatmaps showing differential abundance patterns of 1-(1Z-alkenyl),2-acylglycerophosphocholines [GP0103] (C), 1-alkyl,2-acylglycerophosphocholines [GP0102] (D), and diacylglycerophosphocholines [GP0101] (E). Each row represents an individual lipid species, and each column represents a biological replicate. Color intensity indicates the relative abundance of each lipid species across XH001 and XH001Δ*gdpd*_Xhp1_ monocultures. Deletion of *gdpd*_Xhp1_ resulted in distinct phosphatidylcholine remodeling, characterized by increased ether-linked phosphatidylcholine species (GP0102 and GP0103) and reduced diacylglycerophosphocholine species (GP0101), indicating altered phospholipid homeostasis and membrane lipid composition.

**Supplementary Figure 5:**
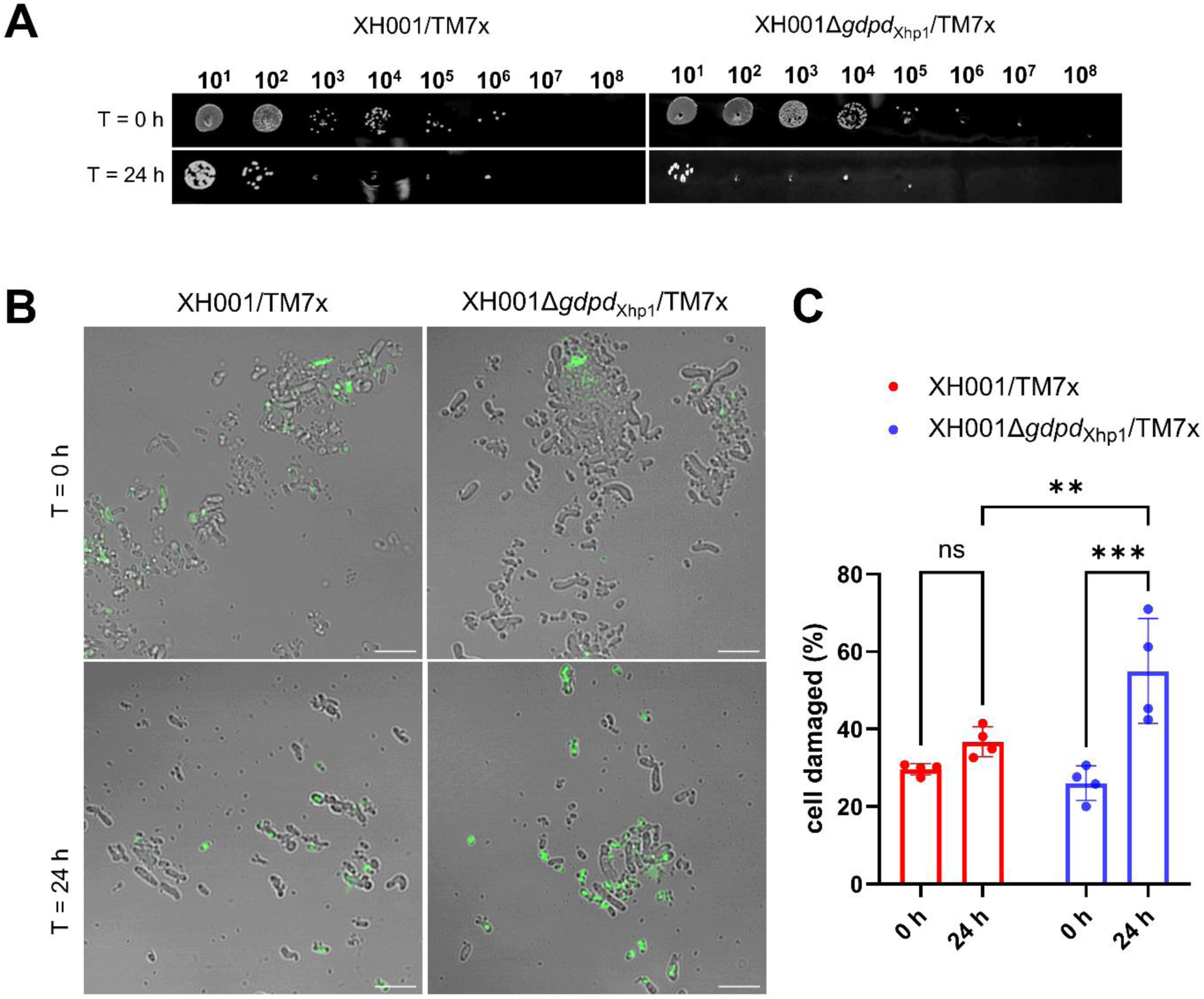
Loss of GDPDₓₕₚ₁ reduces survival and increases cell damage during nutrient starvation. (A) Representative spot-dilution assay of XH001/TM7x and XH001Δ*gdpd*_Xhp1_/TM7x cocultures immediately after transfer to PBS (T = 0 h) and following 24 h incubation in nutrient-free PBS (T = 24 h). The Δ*gdpd*_Xhp1_/TM7x coculture exhibited reduced recovery after 24 h compared with the WT coculture. (B) Representative Sytox Green fluorescence images of XH001/TM7x and XH001Δ*gdpd*_Xhp1_/TM7x cocultures at T = 0 h and T = 24 h. Sytox Green-positive cells indicate membrane-compromised or damaged cells. Scale bars, 5 μm. (C) Quantification of Sytox Green-positive cells shown as the percentage of damaged cells at T = 0 h and T = 24 h. Statistical significance was determined using an unpaired two-tailed Student’s *t*-test. \*\**P* <0.01; \*\*\**P* <0.001, ns = not significant.

**Supplementary Figure 6:**
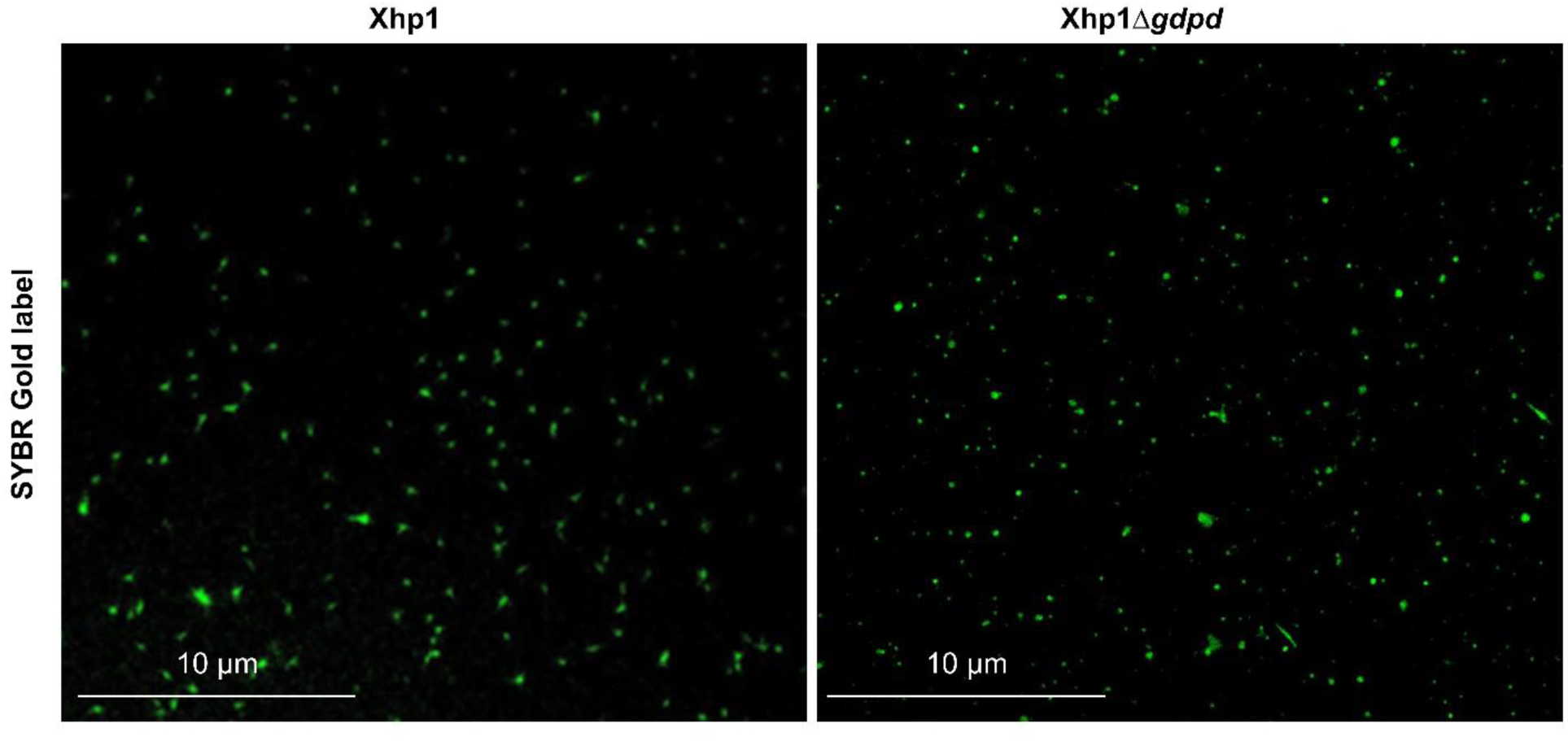
Visualization of isolated Xhp1 and Xhp1Δ*gdpd* phage particles by SYBR Gold staining. Prophages were induced from mid-log-phase cultures of *S. odontolytica* XH001 (OD₆₀₀ ≈ 0.5) with mitomycin C (0.01 μg mL⁻¹) for 3 h. Phage particles released from the induced cultures were recovered and purified, stained with SYBR Gold, and visualized by fluorescence microscopy. Representative images of WT Xhp1 (left) and Xhp1Δ*gdpd* (right) preparations are shown. Scale bars, 10 μm.

**Supplementary Table 1:**
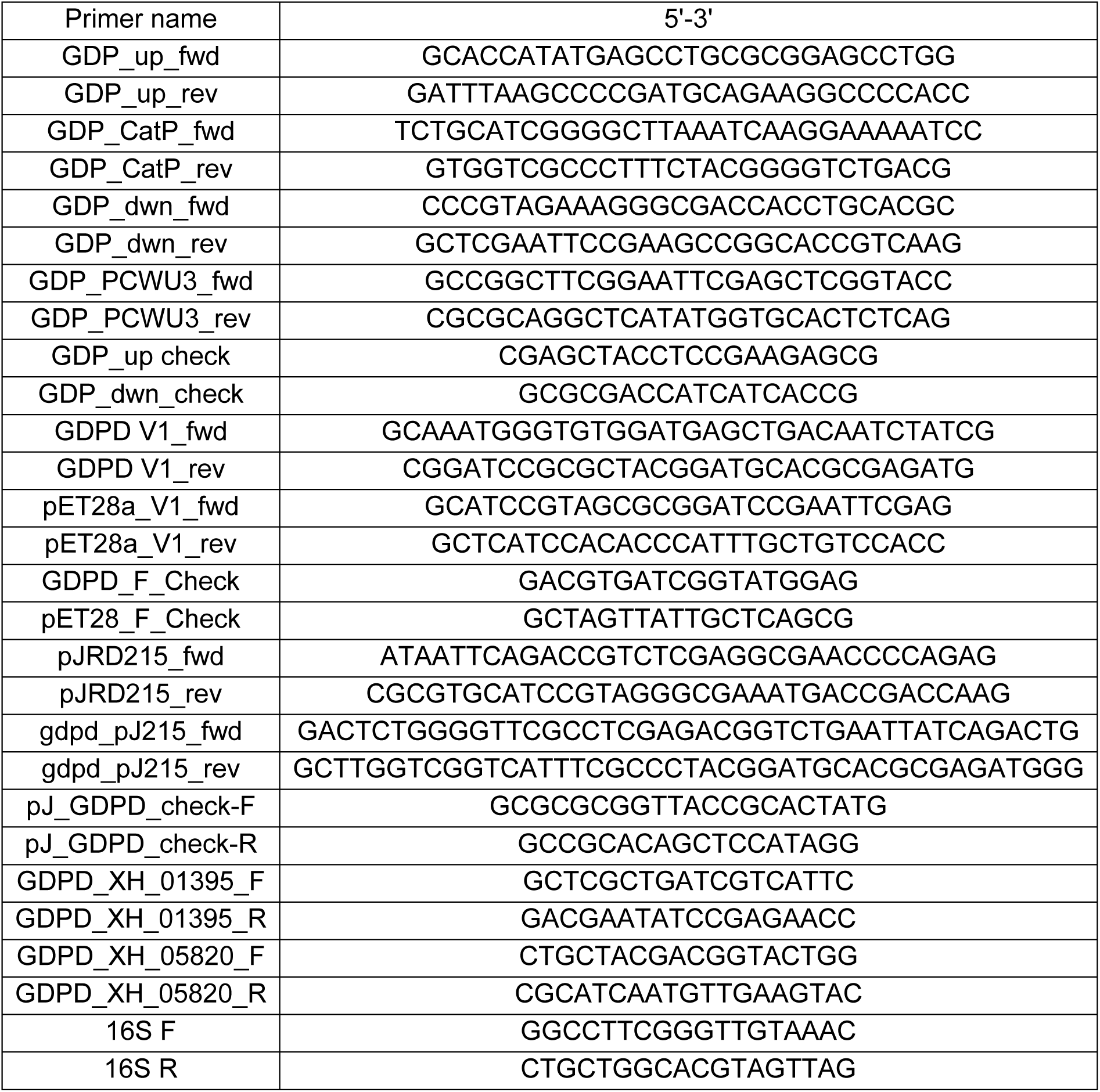
List of primers.

## References

1. Gamachu, S. & Debalo, M. Review of bacteriophage and its applications. Int. J. Vet. Sci. Res 8, 133–147 (2022).

2. Canchaya, C., Fournous, G. & Brüssow, H. The impact of prophages on bacterial chromosomes. Mol Microbiol 53, 9–18 (2004).

3. Bobay, L.M., Touchon, M. & Rocha, E.P. Pervasive domestication of defective prophages by bacteria. Proc Natl Acad Sci U S A 111, 12127–12132 (2014).

4. Wendling, C.C., Refardt, D. & Hall, A.R. Fitness benefits to bacteria of carrying prophages and prophage-encoded antibiotic-resistance genes peak in different environments. Evolution 75, 515–528 (2021).

5. Bucher, M.J. & Czyż̇, D.M. Phage against the machine: the SIE-ence of superinfection exclusion. Viruses 16 (2024).

6. Lang, A.S., Zhaxybayeva, O. & Beatty, J.T. Gene transfer agents: phage-like elements of genetic exchange. Nature Reviews Microbiology 10, 472–482 (2012).

7. Leiman, P.G. et al. Type VI secretion apparatus and phage tail-associated protein complexes share a common evolutionary origin. Proc Natl Acad Sci U S A 106, 4154–4159 (2009).

8. He, X. et al. Cultivation of a human-associated TM7 phylotype reveals a reduced genome and epibiotic parasitic lifestyle. Proceedings of the National Academy of Sciences 112, 244–249 (2015).

9. Bor, B. et al. Phenotypic and physiological characterization of the epibiotic interaction between TM7x and its basibiont Actinomyces. Microbial ecology 71, 243–255 (2016).

10. McLean, J.S. et al. Draft genome sequence of *Actinomyces odontolyticus* subsp. actinosynbacter strain XH001, the basibiont of an oral TM7 epibiont. Genome announcements 4, 10.1128/genomea. 01685–01615 (2016).

11. Bor, B. et al. Rapid evolution of decreased host susceptibility drives a stable relationship between ultrasmall parasite TM7x and its bacterial host. Proceedings of the National Academy of Sciences 115, 12277–12282 (2018).

12. Bor, B., Bedree, J.K., Shi, W., McLean, J.S. & He, X. Saccharibacteria (TM7) in the human oral microbiome. J Dent Res 98, 500–509 (2019).

13. Hendrickson, E.L. et al. Ultrasmall epibiont *Nanosynbacter lyticus* strain TM7x and host bacteria transcriptional activity after initial host parasitism. J Oral Microbiol 16, 2287349 (2024).

14. Hendrickson, E.L. et al. Transcriptome of epibiont Saccharibacteria *Nanosynbacter lyticus* strain TM7x during the establishment of symbiosis. J Bacteriol 204, e0011222 (2022).

15. Dong, P.-T. et al. Episymbiotic *Saccharibacteria* induce intracellular lipid droplet production in their host bacteri. The ISME Journal 18 (2024).

16. Hejazian, S.M., Pirmoradi, S., Zununi Vahed, S., Kumar Roy, R. & Hosseiniyan Khatibi, S.M. An update on glycerophosphodiester phosphodiesterases; from bacteria to human. The Protein Journal 43, 187–199 (2024).

17. Zhang, Y.-M. & Rock, C.O. Membrane lipid homeostasis in bacteria. Nature Reviews Microbiology 6, 222–233 (2008).

18. Parsons, J.B. & Rock, C.O. Bacterial lipids: metabolism and membrane homeostasis. Prog Lipid Res 52, 249–276 (2013).

19. Wittmann, J., Gartemann, K.-H., Eichenlaub, R. & Dreiseikelmann, B. Genomic and molecular analysis of phage CMP1 from *Clavibacter michiganensis* subspecies michiganensis. Bacteriophage 1, 6–14 (2011).

20. Kwan, T., Liu, J., DuBow, M., Gros, P. & Pelletier, J. The complete genomes and proteomes of 27 *Staphylococcus aureus* bacteriophages. Proc Natl Acad Sci U S A 102, 5174–5179 (2005).

21. Vandersteegen, K. et al. Microbiological and molecular assessment of bacteriophage ISP for the control of *Staphylococcus aureus*. PLoS One 6, e24418 (2011).

22. Gu, J. et al. Complete genome sequence of *Staphylococcus aureus* bacteriophage GH15. J Virol 86, 8914–8915 (2012).

23. Cui, Z. et al. Complete genome sequence of wide-host-range *Staphylococcus aureus* phage JD007. J Virol 86, 13880–13881 (2012).

24. Wang, S., Kong, J., Gao, C., Guo, T. & Liu, X. Isolation and characterization of a novel virulent phage (phiLdb) of *Lactobacillus delbrueckii*. Int J Food Microbiol 137, 22–27 (2010).

25. Riipinen, K.A., Forsman, P. & Alatossava, T. The genomes and comparative genomics of *Lactobacillus delbrueckii* phages. Arch Virol 156, 1217–1233 (2011).

26. Casey, E. et al. Molecular characterization of three *Lactobacillus delbrueckii* subsp. bulgaricus phages. Appl Environ Microbiol 80, 5623–5635 (2014).

27. Dupont, K., Janzen, T., Vogensen, F.K., Josephsen, J. & Stuer-Lauridsen, B. Identification of *Lactococcus lactis* genes required for bacteriophage adsorption. Appl Environ Microbiol 70, 5825–5832 (2004).

28. Mahony, J. et al. Investigation of the relationship between lactococcal host cell wall polysaccharide genotype and 936 phage receptor binding protein phylogeny. Appl Environ Microbiol 79, 4385–4392 (2013).

29. Gill, J.J. Revised genome Sequence of *Staphylococcus aureus* bacteriophage K. Genome Announc 2 (2014).

30. Cornelissen, A. et al. The baseplate of *Lactobacillus delbrueckii* bacteriophage Ld17 harbors a glycerophosphodiesterase. J Biol Chem 291, 16816–16827 (2016).

31. Shen, M. et al. A Linear plasmid-Like prophage of *Actinomyces odontolyticus* promotes biofilm assembly. Appl Environ Microbiol 84 (2018).

32. Kumar, A. et al. A host-encoded prophage targets a Candidate Phyla Radiation bacterium and shapes episymbiotic interactions. Proc Natl Acad Sci U S A 123, e2615890123 (2026).

33. Lin, H. et al. Microsecond fingerprint stimulated Raman spectroscopic imaging by ultrafast tuning and spatial-spectral learning. Nature Communications 12, 3052 (2021).

34. Ohshima, N. et al. *Escherichia coli* cytosolic glycerophosphodiester phosphodiesterase (UgpQ) requires Mg^2+^, Co^2+^, or Mn^2+^ for its enzyme activity. J Bacteriol 190, 1219–1223 (2008).

35. Shi, L., Liu, J.-F., An, X.-M. & Liang, D.-C. Crystal structure of glycerophosphodiester phosphodiesterase (GDPD) from *Thermoanaerobacter tengcongensis*, a metal ion-dependent enzyme: Insight into the catalytic mechanism. *Proteins: Structure*, Function, and Bioinformatics 72, 280–288 (2008).

36. Alvarez, H.M., Lanfranconi, M.P. & Hernández, M.A. Metabolism-lipid droplet-nucleic acid crosstalk to regulate lipid storage and other cellular processes in oleaginous *Rhodococcus* bacteria. Biology of the Cell 117, e2400094 (2025).

37. Lee, J. & Ridgway, N.D. Substrate channeling in the glycerol-3-phosphate pathway regulates the synthesis, storage and secretion of glycerolipids. Biochim Biophys Acta Mol Cell Biol Lipids 1865, 158438 (2020).

38. Hishikawa, D., Hashidate, T., Shimizu, T. & Shindou, H. Diversity and function of membrane glycerophospholipids generated by the remodeling pathway in mammalian cells. J Lipid Res 55, 799–807 (2014).

39. Parasassi, T., De Stasio, G., d’Ubaldo, A. & Gratton, E. Phase fluctuation in phospholipid membranes revealed by Laurdan fluorescence. Biophys J 57, 1179–1186 (1990).

40. Rangholia, N., Leisner, T.M. & Holly, S.P. Bioactive ether lipids: primordial modulators of cellular signaling. Metabolites 11 (2021).

41. Bergan, J. et al. The ether lipid precursor hexadecylglycerol protects against Shiga toxins. Cell Mol Life Sci 71, 4285–4300 (2014).

42. Müller, A. et al. Daptomycin inhibits cell envelope synthesis by interfering with fluid membrane microdomains. Proc Natl Acad Sci U S A 113, E7077–e7086 (2016).

43. Grossman, A.S. et al. *Saccharibacteria* deploy two distinct type IV pili, driving episymbiosis, host competition, and twitching motility. The ISME Journal 19 (2025).

44. Iglesias, S.M., Li, F., Briani, F. & Cingolani, G. Viral genome delivery across bacterial cell surfaces. Annu Rev Microbiol 78, 125–145 (2024).

45. Potekhina, N.y.V., Streshinskaya, G.M., Tul’skaya, E.M. & Shashkov, A.S. in Methods in Microbiology, Vol. 38. (eds. F. Rainey & A. Oren) 131–164 (Academic Press, 2011).

46. Shiraishi, T., Yokota, S., Fukiya, S. & Yokota, A. Structural diversity and biological significance of lipoteichoic acid in Gram-positive bacteria: focusing on beneficial probiotic lactic acid bacteria. Biosci Microbiota Food Health 35, 147–161 (2016).

47. Brown, S., Santa Maria, J.P., Jr. & Walker, S. Wall teichoic acids of gram-positive bacteria. Annu Rev Microbiol 67, 313–336 (2013).

48. Zhong, Q. et al. Episymbiotic Saccharibacteria TM7x modulates the susceptibility of its host bacteria to phage infection and promotes their coexistence. Proceedings of the National Academy of Sciences 121, e2319790121 (2024).

49. Brenes, L.R. & Laub, M.T. *E. coli* prophages encode an arsenal of defense systems to protect against temperate phages. Cell Host Microbe 33, 1004–1018.e1005 (2025).

50. Du Toit, A. Prophage against phage. Nature Reviews Microbiology 23, 681–681 (2025).

51. Patel, P.H. & Maxwell, K.L. Prophages provide a rich source of antiphage defense systems. Current Opinion in Microbiology 73, 102321 (2023).

52. Hachmann, A.B. et al. Reduction in membrane phosphatidylglycerol content leads to daptomycin resistance in *Bacillus subtilis*. Antimicrob Agents Chemother 55, 4326–4337 (2011).

53. Sohlenkamp, C. & Geiger, O. Bacterial membrane lipids: diversity in structures and pathways. FEMS Microbiol Rev 40, 133–159 (2016).

54. Fernandes, S. & São-José, C. Enzymes and mechanisms employed by tailed bacteriophages to breach the bacterial cell barriers. Viruses 10 (2018).

55. Knecht, L.E., Veljkovic, M. & Fieseler, L. Diversity and function of phage encoded depolymerases. Frontiers in Microbiology **Volume** 10 - 2019 (2020).

56. Nahar, N. et al. Ultrasmall episymbiont *Nanosynbacter lyticus* employs multiple ATP-generating metabolic pathways during horizontal transmission. The ISME Journal 20 (2025).

57. Bedree, J.K. et al. Quorum sensing modulates the epibiotic-parasitic relationship between *Actinomyces odontolyticus* and its Saccharibacteria epibiont, a *Nanosynbacter lyticus* Strain, TM7x. Front Microbiol 9, 2049 (2018).

58. Bedree, J.K. et al. Identifying essential genes in *Schaalia odontolytica* using a saturated transposon library. J Bacteriol 207, e0016425 (2025).

59. Wu, C. et al. Lethality of sortase depletion in *Actinomyces oris* caused by excessive membrane accumulation of a surface glycoprotein. Mol Microbiol 94, 1227–1241 (2014).

60. Bibek, G.C., Zhou, P., Naha, A., Gu, J. & Wu, C. Development of a xylose-inducible promoter and riboswitch combination system for manipulating gene expression in *Fusobacterium nucleatum*. Applied and Environmental Microbiology 89, e00667–00623 (2023).

61. Wu, C. & Ton-That, H. Allelic exchange in *Actinomyces oris* with mCherry fluorescence counterselection. Applied and Environmental Microbiology 76, 5987–5989 (2010).

62. Cen, L. et al. Exploitation of a bacterium-encoded lytic transglycosylase by a human oral lytic phage to facilitate infection. J Virol 96, e0106322 (2022).

63. Lu, S. et al. Genomic and proteomic analyses of the terminally redundant genome of the Pseudomonas aeruginosa phage PaP1: establishment of genus PaP1-like phages. PLoS One 8, e62933 (2013).

64. Kropinski, A.M., Mazzocco, A., Waddell, T.E., Lingohr, E. & Johnson, R.P. Enumeration of bacteriophages by double agar overlay plaque assay. Methods Mol Biol 501, 69–76 (2009).

65. Raphael Hans, L. Agar overlay (double-layer agar) technique for bacteriophage assays. The Phage (2023).

66. Lee, K. et al. Raman microspectroscopy for microbiology. Nature Reviews Methods Primers 1 (2021).

67. Love, M.I., Anders, S., Kim, V. & Huber, W. RNA-Seq workflow: gene-level exploratory analysis and differential expression. F1000Res 4, 1070 (2015).

68. Priyadarshini, R., Popham David, L. & Young Kevin, D. Daughter cell separation by penicillin-binding proteins and peptidoglycan amidases in *Escherichia coli*. Journal of Bacteriology 188, 5345–5355 (2006).

69. Zamakhaeva, S. et al. Modification of cell wall polysaccharide guides cell division in *Streptococcus mutans*. Nat Chem Biol 17, 878–887 (2021).

